# Cohesin acts as a transcriptional gatekeeper by restraining pause–release to promote processive elongation

**DOI:** 10.1101/2025.09.30.679672

**Authors:** Shoin Tei, Masashige Bando, Toyonori Sakata, Atsunori Yoshimura, Toyoaki Natsume, Masato T Kanemaki, Takashi Sutani, Katsuhiko Shirahige

**Affiliations:** Laboratory of Genome Structure and Function, Institute for Quantitative Biosciences, The University of Tokyo, 1-1-1 Yayoi, Bunkyo-Ku, Tokyo, 113-0032, Japan; Karolinska Institutet, Department of Cell and Molecular Biology, Biomedicum, Tomtebodavägen 16, Stockholm 171 77, Sweden; Department of Chromosome Science, National Institute of Genetics, Research Organization of Information and Systems (ROIS), Yata 1111, Mishima, Shizuoka 411-8540, Japan; Graduate Institute for Advanced Studies, SOKENDAI, Yata 1111, Mishima, Shizuoka 411-8540, Japan; Present address: Research Center for Genome & Medical Sciences, Tokyo Metropolitan Institute of Medical Science, Tokyo, Japan; Department of Biological Science, Graduate School of Science, The University of Tokyo, Bunkyo-ku, Tokyo 113-0033, Japan

## Abstract

Cohesin organizes 3D chromatin architecture, including promoter–enhancer loops, yet its loss has surprisingly modest effects on steady-state gene expression. We address this paradox by demonstrating that cohesin acts at different stages of transcription. First, it promotes Pol II promoter recruitment by facilitating enhancer–promoter communication that maintains active promoter chromatin states. Second, it delays pause release by transiently associating with the transcriptional machinery during the pause–release transition. Kinetic modelling suggests that reduced Pol II recruitment and enhanced pause release have compensatory effects, contributing to minimal changes in steady-state gene expression across genes upon cohesin loss. In contrast, cohesin depletion impairs robust transcriptional induction in response to external stimuli. Moreover, cohesin ensures sufficient pausing duration as a quality-control-like step to promote elongation complex assembly and transcription processivity. Here, we show that cohesin regulates multiple steps of transcription, offering mechanistic insight into cohesin-related diseases, including cancers and cohesinopathies.

## Introduction

Cohesin is a conserved ring-shaped SMC complex whose loading onto chromatin is mediated by the NIPBL–MAU2 heterodimer, which also stimulates cohesin’s ATPase activity^1,2^. During cell division, cohesin ensures proper chromosome segregation by holding replicated sister chromatids together from S phase to anaphase. In interphase, cohesin organizes topologically associated domains (TADs) and chromatin loops, and its depletion severely disrupts these higher-order chromatin structures^3,4^. Given these architectural roles, cohesin has been proposed to regulate gene expression by modulating promoter-enhancer communication^5^. However, gene deletion or protein knockdown of cohesin affects the steady-state expression of only a small fraction of genes^3,6^.

Nevertheless, observations from Cornelia de Lange syndrome (CdLS) and CHOPS syndrome suggest a transcriptional role for cohesin. CdLS is a developmental disorder caused by loss-of-function mutations in cohesin subunits and their regulatory factors^7^. CHOPS syndrome, in contrast, results from gain-of-function mutations in AFF4^8^, the scaffold protein of the super elongation complex (SEC) that promotes RNA polymerase II (Pol II) pause release^9,10^. Notably, CdLS and CHOPS syndrome share transcriptional and phenotypic abnormalities, suggesting a common defect in transcriptional regulation. In metazoan transcription, Pol II frequently pauses shortly after initiation, stabilized by pausing factors such as negative elongation complex (NELF) and DRB sensitivity-inducing factor (DSIF)^11^. CDK9-mediated phosphorylation promotes the release of Pol II from pausing and the transition into productive elongation. CDK9 functions within multiple P-TEFb–containing complexes, including SEC. The shared features of these diseases raised the possibility that cohesin also regulates Pol II pause release, emphasizing the importance of analyzing cohesin’s role at distinct stages of the transcription cycle.

We investigated cohesin’s transcriptional function by genome-wide profiling of transcription dynamics, histone modifications, and 3D genome architecture, together with in vitro reconstitution of the transcription machinery. Our analyses reveal that cohesin acts at different stages of transcription: (i) it promotes Pol II promoter recruitment by facilitating enhancer–promoter communication; and (ii) it ensures proper pausing duration by transiently associating with the transcriptional machinery. When cohesin is depleted, reduced Pol II recruitment and enhanced pause release have compensatory effects, contributing to the minimal changes in steady-state gene expression. In contrast, cohesin depletion impairs robust transcriptional induction in response to external stimuli. Moreover, cohesin’s transient engagement with the transcription machinery acts as a quality-control-like step, ensuring sufficient pausing duration to promote processive elongation and prevent premature transcription termination. These insights have important implications for cohesin-related diseases, including cancers and cohesinopathies, and broader principles of transcriptional regulation.

## Results

### Cohesin is a major determinant of 3D chromatin structure

To investigate how cohesin contributes to transcriptional regulation, we acutely depleted RAD21, a cohesin subunit, in HCT-116 cells expressing an auxin-inducible degron (AID)-tagged RAD21^3,12^. Western blotting confirmed that 3-hour auxin treatment reduced RAD21 protein levels to ∼6%, and ChIP-qPCR at three RAD21 ChIP-seq peaks showed that cohesin binding was reduced to ∼11% (Supplementary Fig. 1a, b). FACS analysis revealed that this treatment had minimal impact on cell cycle progression (Supplementary Fig. 1c).

To examine how cohesin depletion affects higher-order genome organization, we performed Micro-C and obtained ∼1.99 billion valid contacts (Supplementary Fig. 1d). As previously reported^3^, rapid cohesin depletion altered contact distributions, decreasing short-range (<1 Mb) and relatively enriching long-range contacts (Supplementary Fig. 1e). The number of TADs decreased from 4,537 to 2,199, and chromatin loops from 135,539 to 54,332, with the remaining structures exhibiting reduced interaction strength (Supplementary Fig. 1f-h). Upon cohesin depletion, the proportion of loops lacking cohesin peaks at their anchors increased from 40% to 62%, and their median length shortened from 78 kb to 34 kb (Supplementary Fig. 1f, i). These observations indicate that cohesin depletion preferentially disrupts cohesin-mediated loops, which are typically longer (∼165 kb) than loops without cohesin (∼28 kb) (Supplementary Fig. 1j). Accordingly, cohesin-anchored loops exhibited the most pronounced reduction in contact intensity after depletion (Supplementary Fig. 1k). Together, these results confirm cohesin as a major determinant of 3D genome organization.

### Cohesin loss has limited effects on steady-state gene expression

To assess how acute cohesin depletion affects gene expression, we profiled both mature and nascent transcripts using RNA-seq, TT-seq, and EU-seq. TT-seq and EU-seq detect newly synthesized RNAs through 4-thiouridine or 5-ethynyluridine (EU) metabolic labeling, respectively. Among 3,597 well-expressed, non-overlapping genes longer than 5 kb, we identified a limited number of differentially expressed genes upon cohesin depletion: 135 by TT-seq, 83 by RNA-seq, and 182 by EU-seq (Fig. 1a, Supplementary Fig. 2a, b). Of these, 28 genes were shared across all three assays (Supplementary Fig. 2c). Similarly, RNA-seq in SMC1-depleted cells revealed 85 genes with altered expression, 50 of which overlapped with those identified in RAD21-depleted cells (Supplementary Fig. 2d-g). Moreover, genes downregulated upon RAD21 depletion as identified by TT-seq, were preferentially enriched near super-enhancers, as previously reported (Supplementary Fig. 2h)^3^. We further found that both upregulated and downregulated genes tend to exhibit more tissue-restricted expression compared with unchanged genes (Supplementary Fig. 2i, j). Consistent with previous reports^3,4^, these findings indicate that rapid cohesin depletion has only a modest effect on gene expression.

**Fig. 1.**
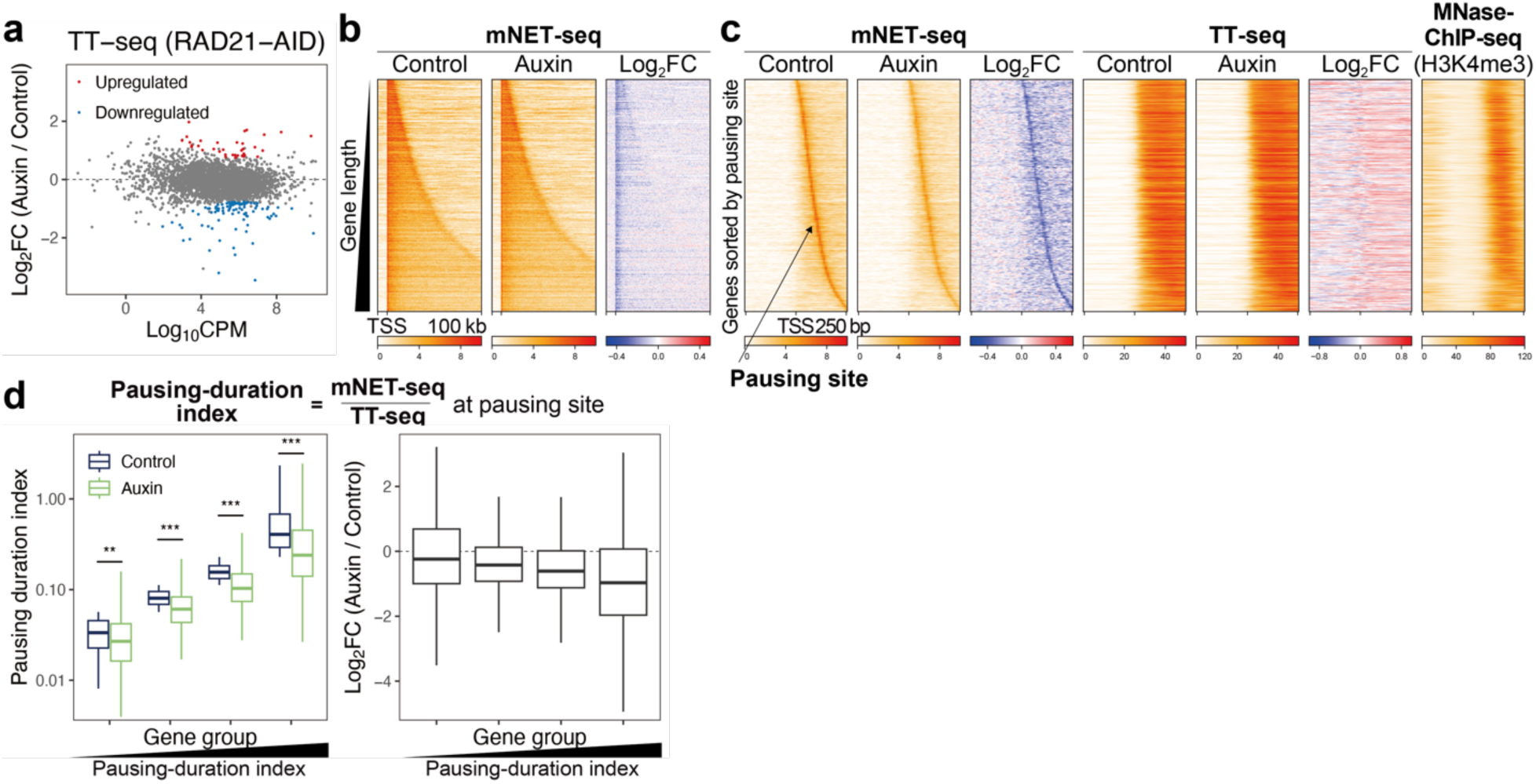
Cohesin promotes Pol II pausing at highly paused genes. **a** MA plot of TT-seq showing gene expression changes upon cohesin depletion. HCT-116 cells expressing AID-tagged RAD21 were treated with auxin for 3 h to degrade RAD21. A total of 3,597 genes included in the analysis had TT-seq TPM ≥ 1, length ≥ 5 kb, and were ≥1 kb away from neighboring genes. Differentially expressed genes were identified using edgeR (FDR < 0.05) based on two biological replicates. **b** Heatmaps showing mNET-seq signals across genes (TSS −10 kb to +100 kb) in control and auxin-treated cells for 3,597 genes. Genes are aligned at their transcription start site (TSS) and sorted by gene length. **c** Heatmaps showing mNET-seq and TT-seq in control and auxin-treated cells, alongside MNase-ChIP-seq (H3K4me3) signals in control cells at promoters (TSS −250 bp to +250 bp) for 3,597 genes. Genes are aligned at their TSSs and sorted by the distance between the TSS and the pausing site. Pausing sites were defined as the highest mNET-seq peak between the TSS and the +1 nucleosome. **d** Box plot showing the pausing-duration index in control and auxin-treated cells and their log_2_ fold changes (auxin/control). The pausing-duration index is calculated as the ratio of mNET-seq to TT-seq within ±25 bp of the pausing sites. The 3,597 genes were evenly divided into four groups based on their pausing-duration index in control cells. Statistical significance was calculated using a two-sided Wilcoxon signed-rank test. (**p < 0.01, ***p < 0.001). Exact p-values from left to right are: 0.0012, 3.55×10^−27^, 5.38×10^−43^, and 2.10×10^−36^.

### Cohesin promotes Pol II pausing at highly paused genes

Given this modest transcriptional impact of cohesin loss, we next examined whether cohesin depletion altered the distribution of Pol II along genes. ChIP-seq for RPB1, the largest subunit of Pol II, revealed a reduction in Pol II occupancy specifically at gene promoters, whereas occupancy along gene bodies remained largely unchanged (Supplementary Fig. 3a). To further investigate this decrease at nucleotide resolution, we performed mNET-seq, which maps transcriptionally engaged Pol II in a strand-specific manner by identifying the last incorporated nucleotide^13^, using an antibody recognizing total Pol II. Consistent with the ChIP-seq results, upon cohesin depletion, mNET-seq signal decreased at promoters but remained unchanged across gene bodies (Fig. 1b). For more detailed analysis, we defined the +1 nucleosome as the first H3K4me3-marked nucleosome downstream of the transcription start site (TSS) and identified the pausing site as the highest mNET-seq peak between the TSS and that nucleosome (Supplementary Fig. 3b).

Using this definition, promoter-focused analysis revealed a specific reduction in mNET-seq signal at pausing sites upon cohesin depletion (Fig. 1c, Supplementary Fig. 3c), while TT-seq remained unchanged, suggesting weakened pausing rather than decreased transcription. To test this quantitatively, we evaluated pausing with a pausing-duration index—the mNET-seq/TT-seq ratio within ±25 bp of the pausing sites (Fig. 1d)^14,15^. This index is theoretically proportional to the dwell time of Pol II at pause sites. Cohesin loss reduced this index (P = 3.67 × 10^−167^, Wilcoxon signed-rank test), with the largest decrease at highly paused genes, indicating shortened pausing.

To confirm the reduction in Pol II pausing following cohesin depletion, we monitored pause release by measuring the Ser2P/RPB1 ratio at promoters, as Ser2 phosphorylation (Ser2P) marks Pol II that has exited the paused state^16^. The Ser2P/RPB1 ratio was negatively correlated with the pausing-duration index, supporting the validity of this measurement, and was increased after cohesin loss, corroborating reduced pausing (Supplementary Fig. 3d, e). As an alternative approach to quantify pausing, we estimated the half-life of paused Pol II by tracking the decay of Pol II promoter binding after triptolide treatment, which blocks new Pol II loading by inhibiting the helicase activity of TFIIH (Supplementary Fig. 3f, g)^17^. Cohesin depletion shortened the average half-life from 14.4 min to 10.3 min, consistent with weakened pausing (Supplementary Fig. 3h). Next, we evaluated Pol II pausing using RPB1 ChIP-seq by calculating a pausing index—promoter occupancy divided by gene-body occupancy (Supplementary Fig. 3i). In line with the pausing-duration index, this index decreased after RAD21 or SMC1 depletion (Supplementary Fig. 3j, k). In contrast, the index increased upon WAPL knockdown, which elevates chromatin-bound cohesin by preventing its dissociation (Supplementary Fig. 3l-n)^18^. We further calculated a pausing index using published PRO-seq data generated in the same cell line and found that cohesin depletion similarly reduced Pol II pausing (Supplementary Fig. 3o)^3^. Although ChIP-seq–based measurements cannot distinguish strand-specific Pol II, we combined these analyses with strand-specific approaches. Together, these five independent approaches demonstrate that cohesin stabilizes Pol II pausing and that its loss leads to a genome-wide reduction in pausing.

Given the reduced pausing observed upon cohesin depletion, we investigated how cohesin loss affects the pausing complex, which includes paused Pol II and its associated factors. Pausing is stabilized the association of NELF with Pol II, and CDK9-mediated phosphorylation triggers NELF dissociation during pause release. To examine the relationship between NELF and Pol II, we measured promoter occupancy of the NELFCD subunit relative to Pol II. Cohesin depletion decreased NELFCD occupancy, while CDK9 inhibition with Flavopiridol, which captures the pausing state, restored this reduction (Supplementary Fig. 3p, q). This rescue indicates that cohesin depletion does not impair the association between paused Pol II and NELF. Instead, the initial decrease in NELFCD likely reflects shortened pausing duration, as NELF dissociates when Pol II is released from pausing^11^. In contrast, cohesin depletion increased promoter occupancy of CDK9 and AFF4, components of the super elongation complex (SEC) that promote pause release. Unlike NELF, CDK9 and AFF4 remained elevated even when CDK9 activity was inhibited by Flavopiridol or DRB. The sustained increase suggests that cohesin depletion enhanced the association of SEC and Pol II at promoters.

### Cohesin constrains early Pol II elongation

Given that cohesin loss shortens promoter pausing, we next examined its effect on elongating Pol II. We first quantified elongation rate using a velocity index—the TT-seq/mNET-seq ratio across gene bodies^19^. This index reflects RNA output per engaged Pol II, such that higher values indicate faster elongation (Supplementary Fig. 4a). Because elongation rate reflects the distance traveled by Pol II per unit time, this velocity index is the reciprocal of the pausing-duration index. Analysis in 1-kb bins revealed increased velocity from the TSS to +8 kb upon cohesin depletion (Supplementary Fig. 4b). To assess velocity changes across entire gene, we divided each gene into three equal regions from the TSS to the gene end (5’, Mid, and 3’). Elevated velocity near the TSS gradually returned to baseline toward the 3’ end, especially in longer genes (Supplementary Fig. 4c). To independently confirm accelerated elongation, we analyzed intronic RNA-seq coverage slopes, where co-transcriptional splicing causes a 5′ to 3′ decline in coverage^20^. In both RAD21- and SMC1-deleted cells, introns within 5 kb of the TSS showed shallower slopes, indicating faster elongation, whereas downstream introns did not (Supplementary Fig. 4d). Together, these two independent assays demonstrate that cohesin depletion accelerates Pol II progression immediately downstream of the TSS.

### Cohesin stabilizes transcription elongation to prevent premature termination

We next asked whether cohesin depletion alters the composition of the elongation complex. Elongation factors, including the PAF1 complex, SPT6, FACT, and DSIF, facilitate processive and stable transcription by interacting with elongating Pol II^11,21^. To quantify these associations, we performed ChIP-seq for each factor and measured their occupancy relative to Pol II across gene bodies. This analysis showed that cohesin loss reduced the relative occupancy of the PAF1 complex (PAF1 and CDC73), SPT6, FACT (SSRP1), and DSIF (SPT4) (Fig. 2a, Supplementary Fig. 4e). In the case of DSIF, which interacts with Pol II from the paused state, we also observed reduced DSIF occupancy around promoters upon cohesin depletion, consistent with shortened pausing. Moreover, co-immunoprecipitation with antibodies against Pol II CTD and Ser2P further revealed that cohesin depletion weakened Pol II interaction with PAF1 components (PAF1, CDC73, or CTR9) by ∼55–75% (Fig. 2b). Because PAF1 associates with Pol II only after pause release^22^, these associations were undetectable during DRB treatment, which blocks Pol II release, but appeared after DRB wash-out. Together, these findings indicate that cohesin promotes the maturation of the elongation complex.

**Fig. 2.**
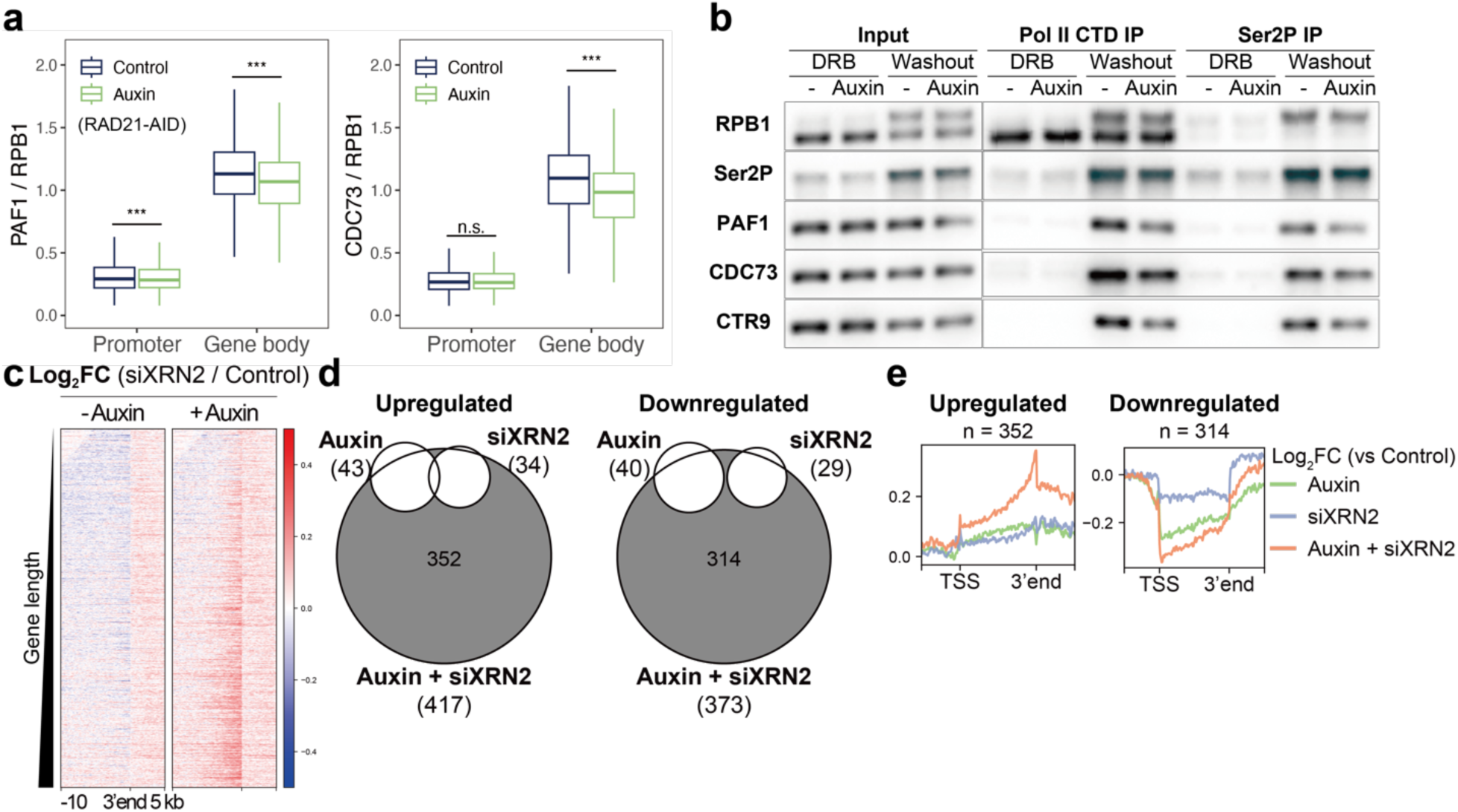
Cohesin stabilizes the elongation complex to suppress premature transcription termination. **a** Box plots showing the relative levels of PAF1 and CDC73 to RPB1 at promoters (TSS −1 kb to +1 kb) and gene bodies (TSS +1 kb to the 3’ gene end) for 3,597 genes in control and auxin-treated cells. Statistical significance was calculated using a two-sided Wilcoxon signed-rank test. (***p < 0.001). Exact p-values: 5.60×10^−6^ (PAF1 promoter), 4.71×10^−121^ (PAF1 gene body), 0.191 (CDC73 promoter), and 1.34×10^−298^ (CDC73 gene body). **b** Co-immunoprecipitation of Pol II and PAF1 complex. Cells were treated with DRB for 3 h and harvested 20 min after DRB removal. Pol II was immunoprecipitated using antibodies against RPB1 CTD or Ser2-phosphorylated RPB1. In the RPB1 IP fraction, PAF1, CDC73, and CTR9 levels normalized to RPB1, were reduced to 59%, 55%, and 75% after auxin treatment. In the Ser2P IP fraction, corresponding levels normalized to Ser2P were reduced to 69%, 73%, and 72%. **c** Heatmaps of log₂ fold changes (siXRN2/control and auxin + siXRN2/auxin) in RNA-seq at gene ends (the 3’ gene end −10 kb to +5 kb) for 3,597 genes. Genes are aligned at their 3’ gene end and sorted by gene length. **d** Venn diagrams showing the overlaps among upregulated or downregulated genes upon auxin, siXRN2, and auxin + siXRN2 treatments. Differentially expressed genes were identified using edgeR (FDR < 0.01) based on three biological replicates. **e** Average profiles of RNA-seq log₂ fold changes (auxin vs control, siXRN2 vs control, and auxin + siXRN2 vs control) for genes uniquely upregulated or downregulated in auxin + siXRN2-treated cells.

Given the incomplete assembly of the elongation complex upon cohesin depletion, we hypothesized that this would promote early transcription termination by impairing transcriptional processivity. To test this, we performed RNA-seq following co-depletion of cohesin and XRN2, a 5’-3’ exoribonuclease that degrades prematurely terminated transcripts (Supplementary Fig. 4f, g)^23^. As expected from its role in transcription termination, XRN2 depletion alone modestly increased signals downstream of 3′ gene ends, with a slightly greater enhancement observed in the absence of cohesin (Fig. 2c, Supplementary Fig. 4h). In contrast, regions upstream of the 3′ ends showed increased RNA-seq signal upon XRN2 knockdown, particularly when cohesin was depleted. This upstream accumulation was more pronounced at longer genes. To further assess transcriptomic changes, we identified genes whose expression was significantly altered in each condition. A total of 666 genes were differentially expressed only upon combined depletion, exceeding the total number of affected genes from either single condition (Fig. 2d). At these synergistically upregulated genes, expression levels exceeded the additive effects of individual depletions (Fig. 2e). Notably, the most prominent increases occurred just upstream of the 3’ ends, consistent with the genome-wide accumulation pattern seen upon co-depletion. In contrast, downregulated genes exhibited expression changes consistent with the combined effects of both treatments. Together, these findings suggest that cohesin depletion leads to the generation of unstable transcripts within genes, likely due to the immature elongation complex, and the XRN2-mediated degradation pathway normally eliminates these transcripts.

### Cohesin and its loader coordinatedly colocalize with RNA Polymerase II at promoters

Cohesin dynamically moves along chromosomes and is enriched at CTCF sites, enhancers, and promoters^24^. The cohesin loader, the NIPBL–MAU2 heterodimer, is required not only for loading cohesin onto DNA but also for the formation of cohesin-mediated loops^25^. Given the observed impact of cohesin depletion on Pol II pausing and elongation, we next investigated how cohesin and its loader are positioned relative to Pol II on chromatin. We performed ChIP-seq to profile RPB1, RAD21, and the loader subunits NIPBL and MAU2 under three conditions: untreated, DRB-treated, and DRB washout. DRB treatment blocks pause release, resulting in Pol II accumulation at promoters (Fig. 3a, Supplementary Fig. 5a). DRB washout relieves the block, allowing Pol II to enter downstream gene bodies, thereby reducing its promoter occupancy (Supplementary Fig. 5b). RAD21, NIPBL, and MAU2 behaved similarly, albeit to a lesser extent, accumulating at promoters during DRB treatment and showing decreased promoter occupancy after washout. These changes indicate that cohesin and its loader associate with promoters in a transcription-dependent manner, similar to Pol II.

**Fig. 3.**
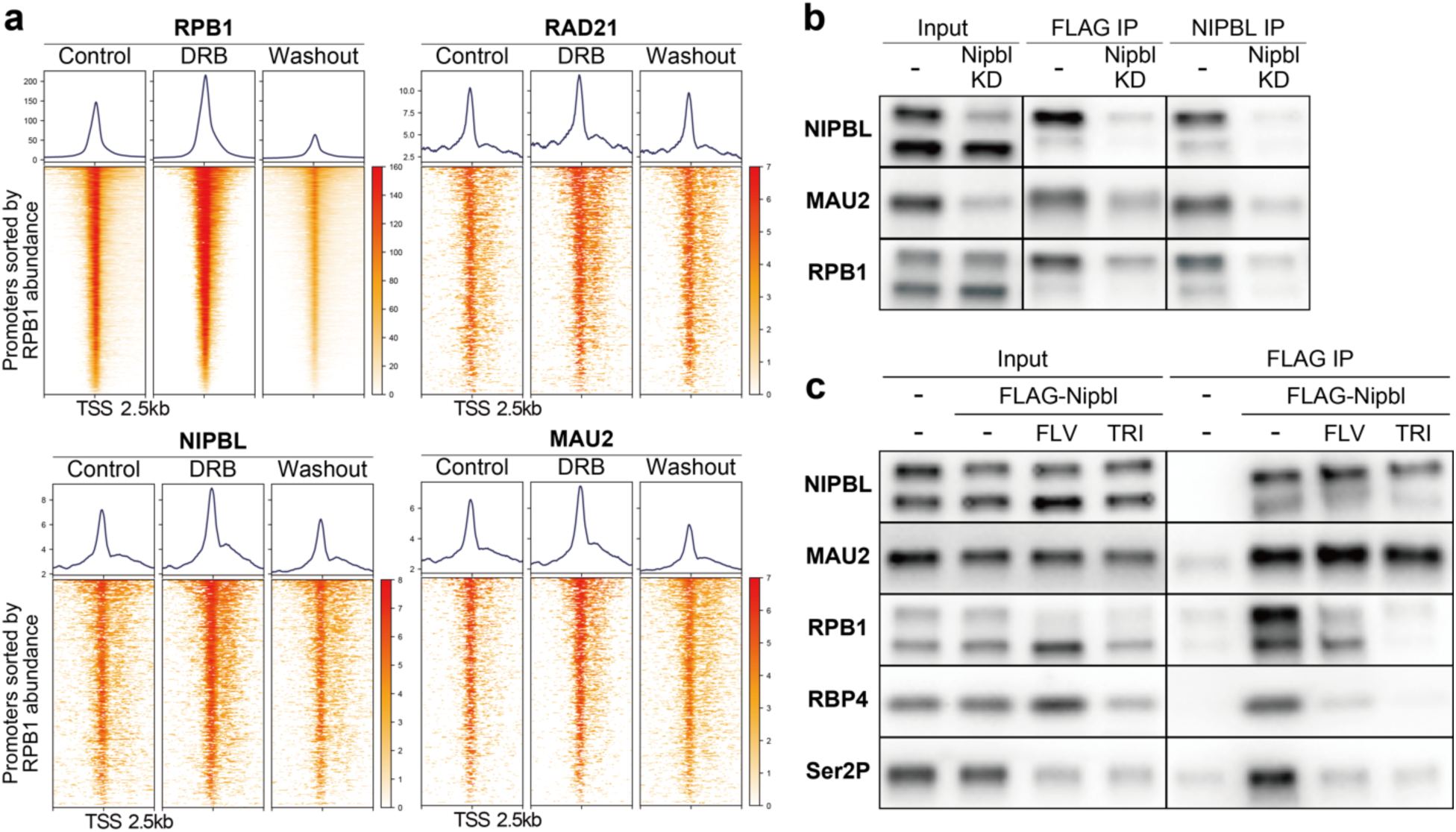
Cohesin and its loader dynamically colocalize with RNA Polymerase II at promoters. **a** Heatmaps of RPB1, RAD21, MAU2, and NIPBL under control, DRB treatment, and DRB washout conditions at promoters (TSS −2.5 kb to +2.5 kb) for 3,597 genes. Promoters are sorted by RPB1 signal intensity in control cells. For the DRB condition, cells were treated with DRB for 3 h. For the washout condition, cells were collected 10 min after DRB removal. **b** Co-immunoprecipitation of the cohesin loader and Pol II. Cells expressing endogenous FLAG-tagged NIPBL were treated with NIPBL-targeting siRNA oligos. NIPBL was immunoprecipitated using antibodies against FLAG or NIPBL. **c** Co-immunoprecipitation of the cohesin loader and Pol II. Cells with or without endogenous FLAG-tagged NIPBL expression were treated with flavopiridol, triptolide, or left untreated. NIPBL was immunoprecipitated using an antibody against FLAG.

We next tested whether the cohesin loader physically associates with Pol II. Co-immunoprecipitation of FLAG-tagged endogenous NIPBL revealed its association with RPB1 and with its loader partner MAU2 (Fig. 3b). TFIIH inhibition by triptolide, which induces degradation of Pol II, abolished the NIPBL-Pol II interaction even though the NIPBL–MAU2 complex remained intact (Fig. 3c). Moreover, the CDK9 inhibitor flavopiridol greatly weakened the NIPBL–Pol II association, indicating that the interaction preferentially occurs downstream of CDK9 activation. In contrast, co-immunoprecipitation using antibodies against cohesin subunits did not detect a clear interaction with Pol II. Given their colocalization at promoters observed by ChIP–seq, cohesin may indirectly interact with Pol II via its loader. Thus, we concluded that cohesin and its loader coordinatedly colocalize with transcriptionally engaged Pol II at promoters, and that CDK9 activity modulates their association, making the interaction between the loader and Pol II detectable by immunoprecipitation.

### Cohesin and its loader engage the transcription machinery at the pause–release transition

Given their colocalization and physical association with Pol II, we investigated whether cohesin and its loader act as components of the transcriptional machinery. We assembled a transcriptional complex on a DNA template containing GAL4-binding sites and TATA promoter in HeLa nuclear extract (Fig. 4a). Upon activation with GAL4-VP16, we observed the recruitment of the cohesin loader, Mediator complex, general transcription factors, Pol II, and both pausing-associated (SPT5, NELFA) and release-associated (AFF4, CDK9) factors to the template (Fig. 4b). NTP addition induced Pol II Ser5 (Ser5P) and Ser2 phosphorylation (Ser2P) of Pol II, whereas DRB selectively inhibited Ser2P. Although the precise position of Pol II pausing on this template remains undefined, the reconstituted system appears to recapitulate transcription factor recruitment upon gene activation and Pol II phosphorylation dynamics (Supplementary Fig. 6a). Notably, while the loader was present, cohesin itself was not clearly detected on the template. Moreover, we detected RNA production from this DNA template by qPCR (Supplementary Fig. 6b). Using recombinant NIPBL fragments, we found that the N-terminal region of NIPBL is required for loader association upon transcriptional activation (Supplementary Fig. 6c-e). Furthermore, Mediator depletion abolished the recruitment of both the loader and Pol II, suggesting that loader recruitment depends on active transcription rather than direct binding by GAL4–VP16 (Supplementary Fig. 6f).

**Fig. 4.**
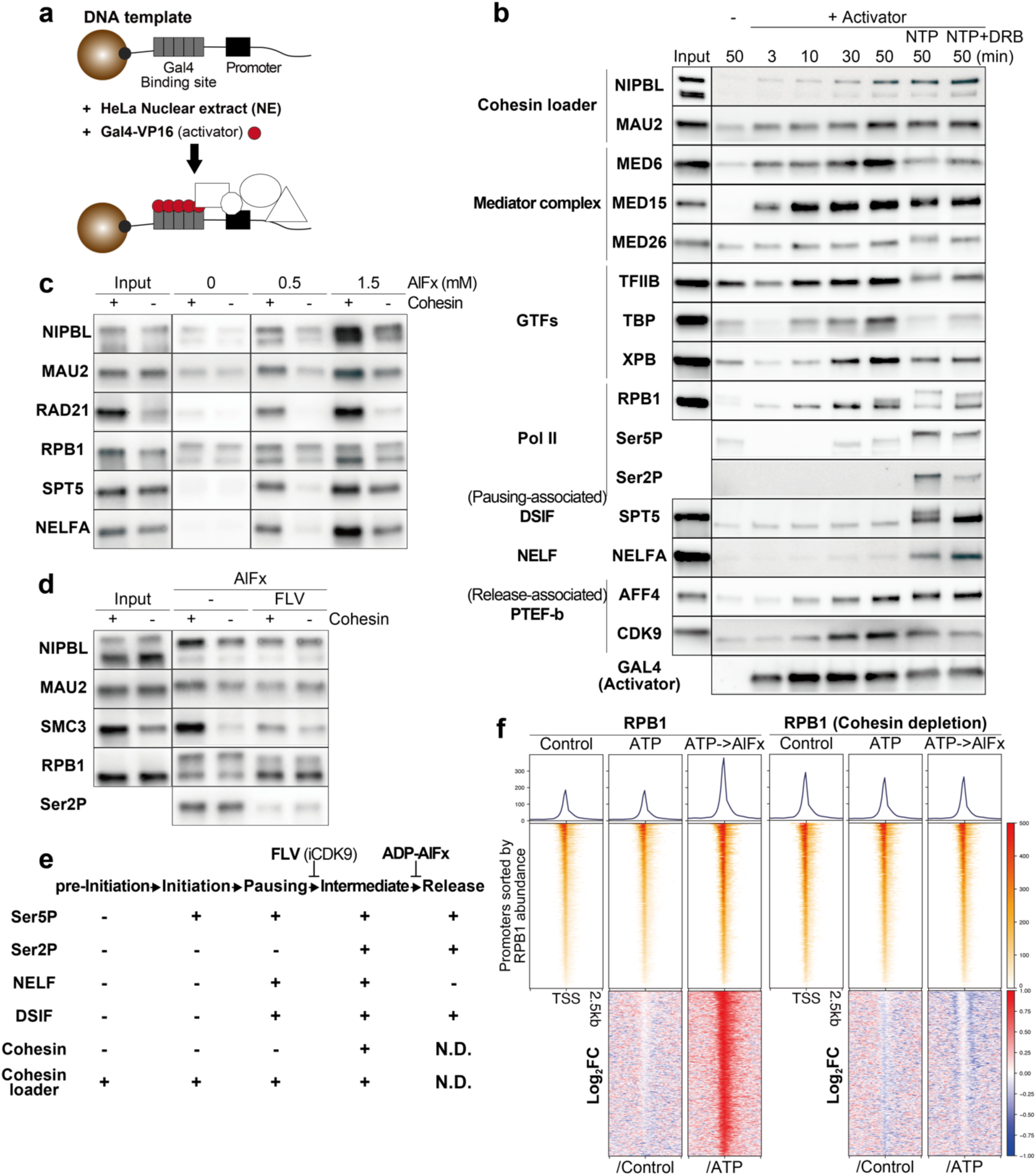
Cohesin and its loader associate with the transcription complex during the pause–release transition. **a** Schematic of the *in vitro* transcription assay. A DNA template containing five GAL4-binding sites and a TATA promoter was preloaded with GAL4-VP16 and incubated with HeLa nuclear extract. After washing, template-bound factors were eluted for analysis. **b** *In vitro* transcription assay to assess factor recruitment upon activator addition. Template DNA preloaded with GAL4-VP16 was incubated with nuclear extracts (Input) for the indicated times. In NTP and NTP + DRB condi-ons, DRB was added at the start of incubation, followed by NTP addition 40 min later. **c** *In vitro* transcription assay with cohesin ATPase inhibition by AlFx. Cohesin was depleted from nuclear extract using antibodies against SMC1/SMC3. A:er a 20 min incuba-on with template DNA and nuclear extract, NTPs were added, followed 3 min later by AlFx (0, 0.5, and 1.5 mM). **d** *In vitro* transcription assay with CDK9 inhibition on AlFx-induced state. Cohesin-depleted nuclear extracts were pre-incubated with CDK9 inhibitor flavopiridol (FLV). NTP and AlFx additions were performed as in c. **e** Schematic summary of factor recruitment across transcriptional stages. The presence or absence of Ser5P, Ser2P, NELF, and DSIF was inferred from previous studies. The cohesin loader was detected upon addition of activator, NTPs, and DRB (b), indicating its presence during the pre-initiation, initiation, and pausing. AlFx-induced state was placed between pausing and release because it contained cohesin, DSIF, and NELF (c) and was abolished by CDK9 inhibition (d). N.D indicates not determined. **f** Heatmaps of RPB1 CUT&Tag signals at promoters (TSS −2.5 kb to +2.5 kb) for 3,597 genes under control, ATP, and ATP->AlFx conditions, with or without cohesin depletion. Permeabilized cells were treated with ATP for 10 min or with ATP for 5 min followed by AlFx for 5 min.

Next, we inhibited cohesin’s ATPase activity with aluminum fluoride (AlFx) to assess its role in transcription^26,27^. AlFx, in complex with ADP, mimics the transition state of ATP hydrolysis, thereby blocking cohesin’s ATPase cycle. AlFx treatment markedly enhanced the recruitment of cohesin, its loader, Pol II, and both pausing- and release-associated factors, making cohesin readily detectable (Supplementary Fig. 6g). This effect was most robust when AlFx was added after ADP, compared to addition before or simultaneous with ADP. Cohesin depletion and CDK9 inhibition abolished the complex (Fig. 4c, d). These findings indicate that: (i) cohesin and its loader are parts of the transcriptional complex, along with pausing- and release-associated factors; (ii) cohesin is required for the assembly of this AlFx-induced complex; and (iii) this complex forms downstream of CDK9 activity, as its formation was enhanced by AlFx addition following CDK9 activation by ADP and abolished by CDK9 inhibition. Collectively, we propose that the AlFx-induced complex represents a transient intermediate state between pausing and pause release, as the NELF complex has been reported to dissociate at the time of Pol II pause release (Fig. 4e). In this model, ATP hydrolysis by cohesin triggers its disassembly as ATPase inhibition with AlFx allowed detection of this otherwise transient state. This transient intermediate may mechanistically explain the shortened Pol II pausing upon cohesin depletion (Fig. 1d). The loss of cohesin likely prevents the formation of this state, thereby promoting premature pause release.

To examine the effects of AlFx in a more physiologically relevant cellular context, we treated cells with AlFx after ATP addition. CUT&Tag revealed enhanced promoter binding of RPB1, RAD21, and NIPBL upon AlFx treatment, which was abolished by cohesin depletion, consistent with our in vitro observations (Fig. 4f, Supplementary Fig. 6h). These results suggest the in vivo relevance of the transient intermediate state between pausing and pause release identified in vitro.

### Cohesin maintains the H3K27ac mark at promoters

Because H3K27ac is a hallmark of active promoters^28^, we investigated whether cohesin depletion altered this modification. ChIP-seq revealed that cohesin loss significantly reduced H3K27ac at 92% of promoters (FDR < 0.05; Fig. 5a, b, Supplementary Fig. 7a). Given that chemical inhibition of H3K27ac is known to diminish promoter binding of the general transcription factor TFIID^29^, we tested whether cohesin loss affects TFIID occupancy. Indeed, promoter occupancy of TFIID subunits TAF1 and TBP decreased at 76% and 91% of promoters, respectively (Fig. 5c, d, Supplementary Fig. 7b, c). As expected from this relationship, promoter H3K27ac levels were strongly correlated with both TAF1 (ρ = 0.77, P < 1 × 10^−300^) and TBP occupancy (ρ = 0.69, P < 1 × 10^−300^), as well as with their respective log₂ fold-changes upon cohesin loss (TAF1: ρ = 0.49, P = 2 × 10^−218^; TBP: ρ = 0.39, P = 5.8 × 10^−134^; Fig. 5e, Supplementary Fig. 7d). These observations suggest that cohesin depletion reduces promoter activity, as indicated by decreased H3K27ac, and consequently impairs TFIID recruitment.

**Fig. 5.**
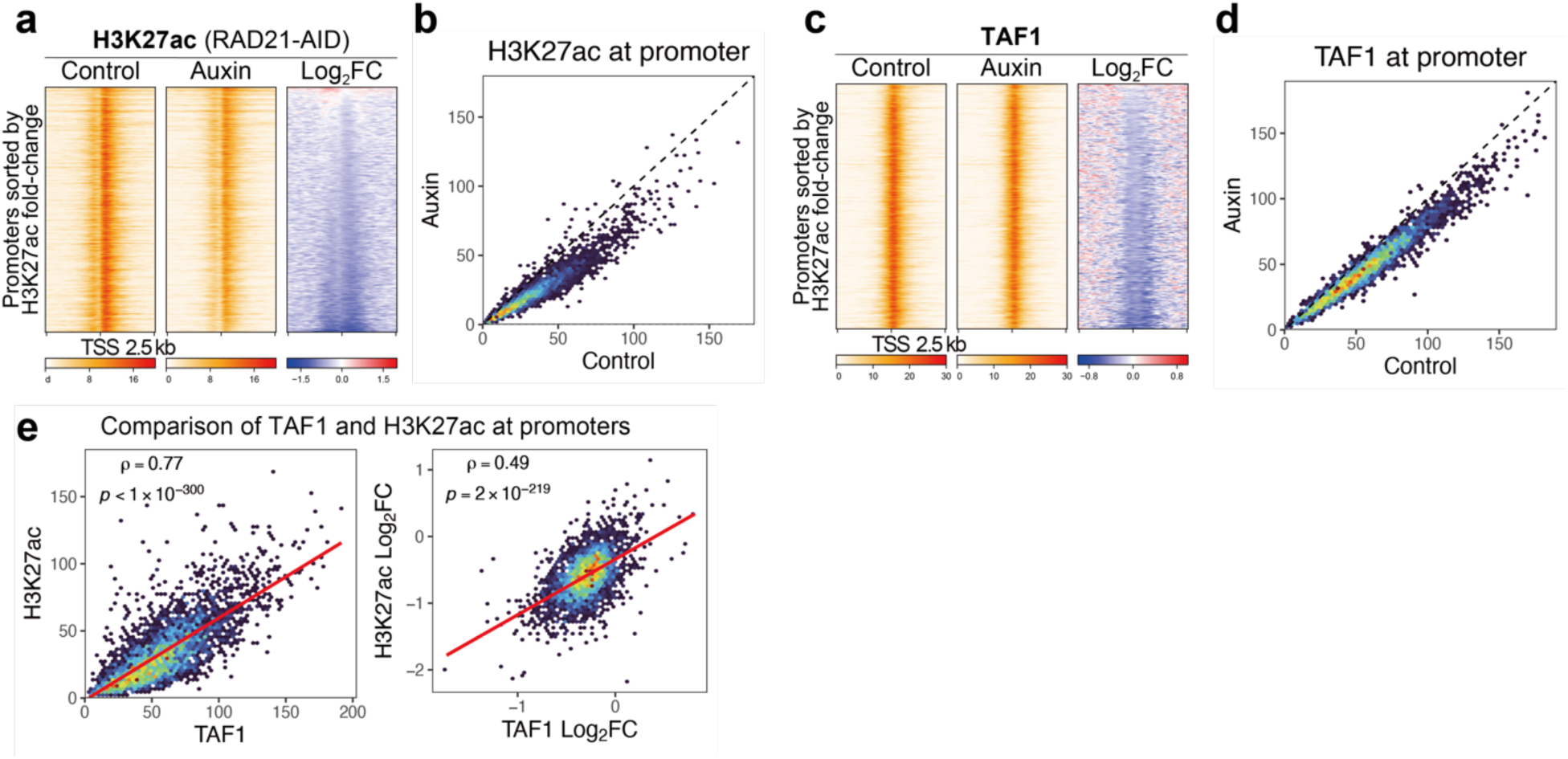
Cohesin maintains the H3K27ac mark at promoters. **a** Heatmaps of H3K27ac ChIP-seq signals at promoters (TSS −2.5 kb to +2.5 kb) for 3,597 genes in control and auxin-treated cells. Promoters are sorted by H3K27ac signal intensity in control cells. **b** Comparison of promoter H3K27ac (TSS −1 kb to +1 kb) between control and auxin-treated cells for 3,597 genes. Upon cohesin depletion, H3K27ac levels decreased at 92% of promoters, increased at 2%, and remained unchanged at 6% (FDR < 0.05, binomial test). **c** Heatmaps of TAF1 ChIP-seq signals at promoters (TSS −2.5 kb to +2.5 kb) for 3,597 genes in control and auxin-treated cells. Promoters are shown in the same order as in (a). **d** Comparison between promoter TAF1 and H3K27ac (TSS −1 kb to +1 kb) in control cells, and their log2 fold changes (auxin/control) for 3,597 genes. P-value (p) and Spearman’s correlation coefficient (ρ) were calculated using Spearman’s rank correlation test.

Because TFIID is required for efficient recruitment of Pol II to promoters, we assessed Pol II recruitment by inhibiting CDK7 with THZ1 to trap Pol II in a promoter-bound, pre-initiation state. Under these conditions, cohesin depletion reduced Pol II occupancy at promoters, which positively correlated with promoter H3K27ac levels (ρ = 0.69, P < 1 × 10^−300^) as well as their log2 fold changes upon cohesin depletion (ρ = 0.39, P < 5.8 × 10^−132^; Supplementary Fig. 7e, f). Together with the largely unchanged Ser5P/RPB1 ratio at promoters (Supplementary Fig. 7g), these results indicate that cohesin loss impairs Pol II recruitment itself rather than initiation-associated phosphorylation. Together, promoter H3K27ac likely reflects Pol II recruitment, given its positive correlation with promoter TFIID occupancy and Pol II levels under CDK7 inhibition.

### Cohesin regulates H3K27ac distribution via promoter–enhancer contacts

Given that cohesin depletion reduces H3K27ac at promoters, we next examined how it affects H3K27ac distribution across both promoters and enhancers, as this modification marks both regulatory elements. We annotated promoters and enhancers genome-wide using ChromHMM^30^. Among 40,142 combined regulatory elements, H3K27ac levels increased at 6,887 sites and decreased at 18,728 upon cohesin depletion (FDR < 0.05; Fig. 6a, b, Supplementary Fig. 8a). Published H3K27ac ChIP-seq data generated in the same cell line also showed similar patterns of gain and loss (Supplementary Fig. 8b). These changes were further validated by ChIP–qPCR at four representative regions, two with increased and two with decreased H3K27ac (Supplementary Fig. 8c). Elements with increased H3K27ac were enriched at H3K4me1-marked enhancers, whereas those exhibiting decreases were primarily localized at H3K4me3-positive promoters (Supplementary Fig. 8d), consistent with the reduction in promoter H3K27ac we observed (Fig. 5a). BRD4, MED1, mNET-seq and TT-seq signals exhibited similar increases and decreases, while H3K4me1/3 levels themselves remained unchanged (Supplementary Fig. 8a, e). Next, we investigated the genomic distribution of H3K27ac-altered elements. Elements with the same H3K27ac response, particularly increased elements, were more often positioned adjacent to each other than expected under a random (uniform) genomic distribution (Fig. 6c). In contrast, increased and decreased sites were located next to each other less often than expected. This positional bias suggests a role for 3D genome architecture in shaping the distribution of H3K27ac.

**Fig. 6.**
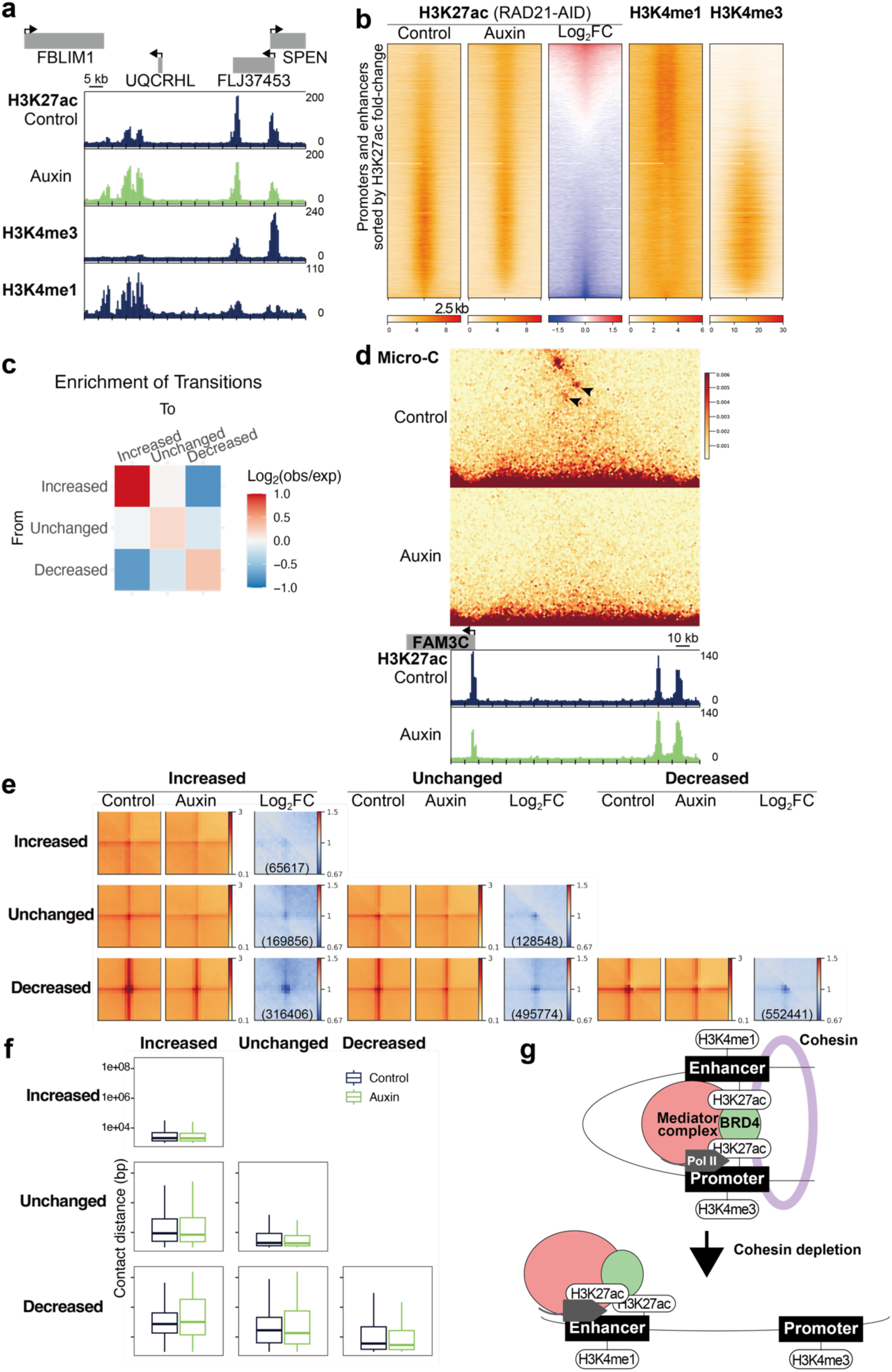
Cohesin regulates the genomic distribution of H3K27ac via promoter–enhancer contacts. **a** Representative genomic region containing regulatory elements with increased or decreased H3K27ac in auxin-treated cells (chr1: 15780000-15860000). H3K4me3 and H3K4me1 tracks are shown to mark promoter and enhancer-associated regions, respectively. **b** Heatmaps of H3K27ac ChIP–seq signals in control and auxin-treated cells, alongside H3K4me1 and H3K4me3 signals in control cells, at 40,142 regulatory elements (promoters and enhancers). Regulatory elements are sorted by fold change (auxin/control) in H3K27ac signal. **c** Heatmap of the log2 fold change (obs/exp) of transitions between consecutive regulatory elements. Regulatory elements were categorized by H3K27ac change upon cohesin depletion (increased, unchanged, or decreased). Transition probabilities were calculated from their order along the genome and compared with the expected probabilities assuming a uniform distribution. **d** Representative genomic region containing regulatory elements with increased or decreased H3K27ac upon auxin treatment (chr7:121380000-121560000). A Micro-C contact matrix is shown to illustrate chromatin contacts between elements with increased and decreased H3K27ac. Black arrowheads indicate these interactions. **e** Aggregate plots of Micro-C contacts between regulatory elements categorized as in c. All six pairwise combinations are shown, with the number of element pairs indicated in parentheses. **f** Box plots of genomic distances of chromatin contacts between the same element pairs as in e. Contacts were defined as paired-end Micro-C reads connecting two regulatory elements. **g** Proposed model illustrating how cohesin-mediated promoter–enhancer interactions support transcription. Cohesin-dependent contacts enable functional communication between enhancers and promoters. Cohesin depletion disrupts this communication, so that enhancers no longer sustain promoter H3K27ac, resulting in reduced H3K27ac at promoters and increased levels at enhancers.

To test this idea, we examined chromatin contacts across the three element classes using Micro-C data. While contacts were observed between all possible pairwise combinations, elements with increased (mostly enhancers) and decreased H3K27ac (mostly promoters) formed the strongest interactions, which were markedly weakened after cohesin depletion (Fig. 6d, e). Analysis of contact distances revealed that interactions between increased and decreased elements spanned the longest genomic distances, both before and after cohesin loss (Fig. 6f). These observations indicate that cohesin is required to maintain long-range promoter–enhancer contacts, consistent with its established role in forming long-range chromatin loops (Supplementary Fig. 1j). We conclude that cohesin loss disrupts promoter–enhancer interactions spanning large genomic distances, resulting in increased H3K27ac at enhancers and decreased H3K27ac at promoters. Moreover, the observed redistribution of H3K27ac between enhancers and promoters suggests that cohesin-dependent contacts enable functional communication between these regulatory elements. In our model, cohesin depletion disrupts this communication, so that enhancers no longer sustain promoter H3K27ac, resulting in reduced H3K27ac at promoters and increased levels at enhancers (Fig. 6g).

### Genes with modest expression changes upon cohesin loss show reduced Pol II recruitment and enhanced pause release

Thus far, our data have shown that cohesin depletion shortens Pol II pausing, accelerates early elongation, and lowers promoter H3K27ac (Supplementary Fig. 9a). However, transcriptome analysis revealed only modest changes in gene expression upon the depletion. To determine how these changes collectively yield modest transcriptional effects, we analyzed their gene-level interplay. We calculated log₂ fold-changes for promoter H3K27ac, pausing-duration index, velocity index, and TT-seq expression, and then applied k-means clustering. To minimize pausing and termination effects, velocity was calculated across the gene body excluding the first and last 1 kb. Unsupervised clustering classified genes into three classes: Cluster 1 (55%) contained genes with little or unchanged expression, while Clusters 2 (23%) and 3 (22%) included genes with overall increased or decreased expression upon cohesin loss, respectively (Fig. 7a, b). Consistently, significantly upregulated genes were mostly found in Cluster 2, and downregulated genes in Cluster 3.

**Fig. 7.**
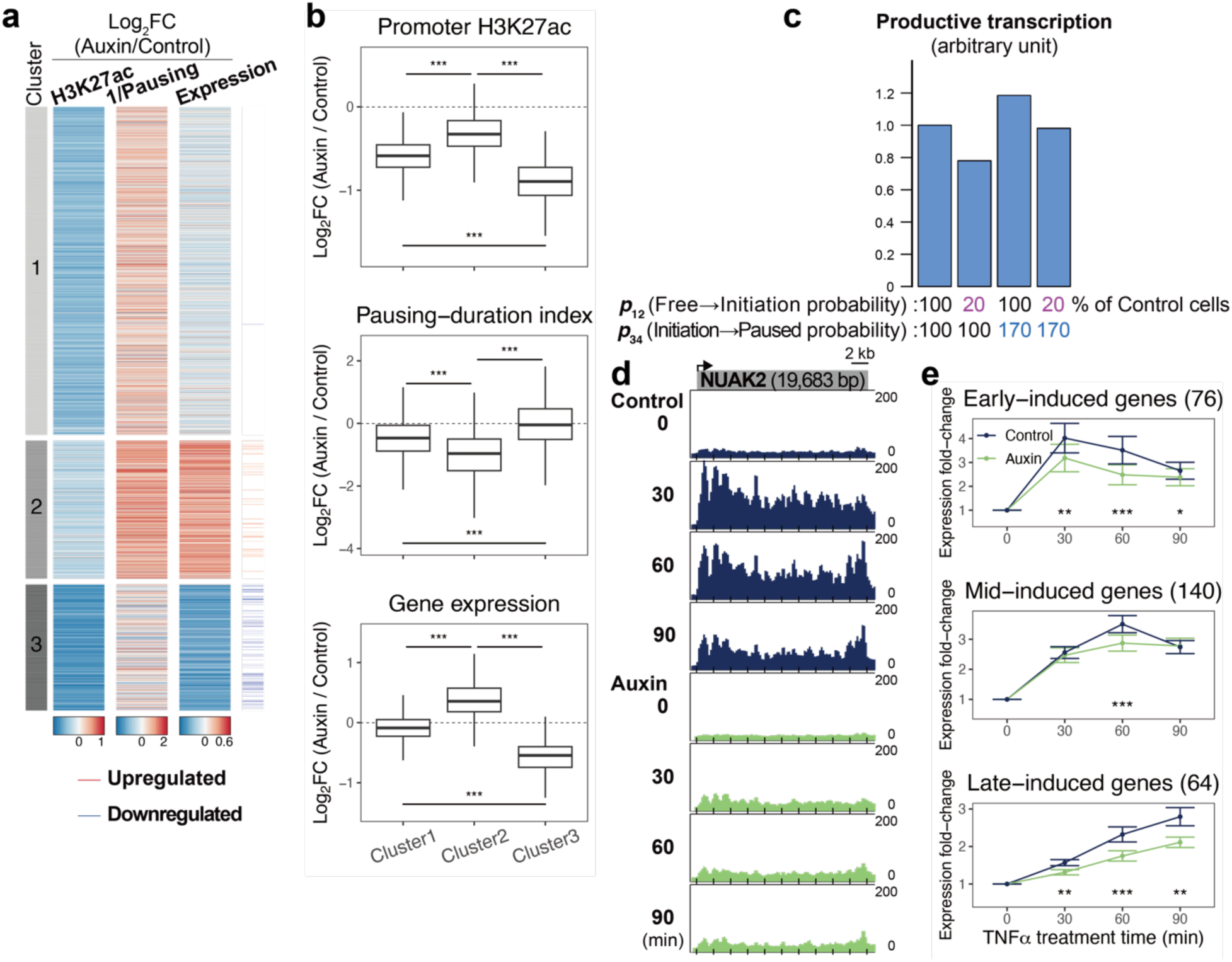
Cohesin depletion reduces initiation but enhances pause release, counterbalancing steady-state gene expression yet impairing rapid gene induction. **a** Heatmaps showing log₂ fold changes (auxin/control) in promoter H3K27ac, pausing-duration index, and TT-seq gene expression. Genes were grouped into three clusters by K-means clustering based on log₂ fold changes in these parameters and the velocity index (Supplementary Fig. 9b). The pausing-duration index was inverted (1/x) to align its directionality with the others. Upregulated or downregulated genes (Fig. 1a) are highlighted with red and blue lines, respectively. **b** Box plots showing log2 fold changes (auxin/control) in promoter H3K27ac, pausing-duration index, and TT-seq gene expression across the clusters in (a). Statistical significance was calculated using pairwise Welch’s t-tests between clusters. (***p < 0.001). Exact p-values for promoter H3K27ac: 3.70×10^−146^ (Cluster1 vs 2), 2.67×10^−158^ (Cluster1 vs 3), 2.33×10-213 (Cluster2 vs 3); for pausing-duration index: 1.21×10^−54^ (Cluster1 vs 2), 1.99×10^−39^ (Cluster1 vs 3), 7.40×10^−96^ (Cluster2 vs 3); for TT-seq gene expression: 8.09×10^−264^ (Cluster1 vs 2), 6.08×10^−264^ (Cluster1 vs 3), 4.26×10^−249^ (Cluster2 vs 3). **c** Bar plots of simulated effects of cohesin depletion on productive transcription. Reduced Pol II recruitment and enhanced pause release were modeled as decreased p_12_ and increased p_34_, respectively. Productive transcription was defined as the probability of Pol II being in the Elongation state and normalized to 1 in the control. **d** Representative gene showing reduced TNFα-induced expression upon auxin treatment. EU-seq was performed at 0, 30, 60, and 90 min after TNFα treatment. **e** Line plots of gene expression fold changes (each time point/ 0 min) at TNFα-induced genes in control and auxin-treated cells. Genes upregulated upon TNFα treatment in control cells (edgeR; FDR < 0.05, fold change > 2) were classified as early, mid, or late-induced genes based on the time point of peak expression, with the number of genes indicated in parentheses. Data represent mean ± standard error. Statistical significance was calculated using a two-sided Wilcoxon rank-sum test. (*p < 0.05, **p < 0.01, ***p < 0.001).

Changes in promoter H3K27ac, Pol II pausing, and velocity also showed distinct patterns across the three clusters. H3K27ac decreased moderately in Cluster 1, minimally in Cluster 2, and most strongly in Cluster 3. Pol II pausing showed the inverse pattern, being slightly reduced in Cluster 1, markedly decreased in Cluster 2, and unchanged in Cluster 3. Velocity was unchanged in Cluster 1, increased in Cluster 2, and decreased in Cluster 3 (Supplementary Fig. 9b, c). These patterns suggest that promoter H3K27ac and Pol II pausing have divergent effects on gene expression, while transcriptional velocity amplifies the effect of pausing. In Cluster 1, comparable reductions in H3K27ac and Pol II pausing act in opposite directions, while elongation velocity remains largely unchanged, resulting in minimal changes in gene expression. In Cluster 2, slight H3K27ac loss, a large reduction in pausing, and increased velocity combine to upregulate gene expression. Conversely, in Cluster 3, pronounced H3K27ac decrease, little change in pausing, and a decline in velocity together suppress transcription. Promoter H3K27ac was used as a proxy for Pol II recruitment to promoters, while reduced Pol II pausing indicates enhanced pause release. Thus, the gene-specific interplay between diminished Pol II recruitment and accelerated pause release likely accounts for the limited impact of cohesin depletion on the steady-state transcriptome. As observed for downregulated genes, genes in Cluster 3 were located closer to super-enhancers than those in the other clusters (Supplementary Fig. 9d). Genes in Cluster 2 and 3 tended to exhibit more tissue-restricted expression compared with Cluster 1 (Supplementary Fig. 9e). Gene Ontology enrichment analysis further revealed that Cluster 1 was enriched for terms related to the mitotic cell cycle and chromosome segregation, whereas Cluster 2 showed enrichment for vesicle-mediated transport and membrane organization (Supplementary Fig. 9f). In contrast, Cluster 3 was preferentially associated with ribonucleoprotein complex biogenesis and DNA damage response.

### Simulations suggest compensatory effects of reduced Pol II recruitment and enhanced pause release

To examine how reduced Pol II recruitment and enhanced pause release might jointly impact transcriptional output, we simulated Pol II transitions through four states: Free, Initiation, Paused, and Elongation (Supplementary Fig. 10a). We then modulated the transition probabilities between these states to reflect changes observed upon cohesin depletion. Transition path and its probability were assigned to reflect previously reported Pol II transitions and their stationary distribution across these states (Supplementary Fig. 10b)^31^. Reducing the Free-to-Initiation probability, recapitulating lower Pol II recruitment observed upon cohesin loss, decreased the Elongation fraction, a proxy for transcriptional output (Supplementary Fig. 10c). In contrast, raising the Paused-to-Elongation probability, reflecting enhanced pause release, increased the Elongation fraction. Notably, when both recruitment and release probabilities were altered together, the proportion of Pol II in Elongation returned to near-baseline (Fig. 7c). These simplified simulations suggest that reduced Pol II recruitment and enhanced pause release have compensatory effects, contributing to largely unchanged gene expression, consistent with our experimental observations. We also asked how these decreases affect the pausing index, designed to mimic the Pol II ChIP-seq pausing index. In the simulation, the pausing index was defined as the ratio of Pol II in the Initiation + Paused states to that in the Elongation state. This index responded specifically to changes in the Paused-to-Elongation probability, supporting the validity of the ChIP–seq–based pausing index (Supplementary Fig. 10d).

### Cohesin loss impairs rapid gene induction upon external stimuli

Because promoter-paused Pol II is thought to serve as a preloaded complex for rapid transcriptional responses to external stimuli^11^, we asked whether cohesin depletion affects cytokine-induced gene activation. EU-seq after TNFα treatment revealed that cohesin depletion reduced gene induction upon TNFα-treatment (Fig. 7d, e, and Supplementary Fig. 11a). Before stimulation, these genes already exhibited lower pausing indices and decreased promoter H3K27ac and TAF1 (Supplementary Fig. 11b-e), suggesting that decreased pausing and/or lowered promoter H3K27ac underlie the impaired induction. These findings suggest that cohesin is required for a robust transcriptional activation in response to external stimuli, indicating its crucial roles in stimulus-induced gene regulation.

## Discussion

Here, we demonstrate that cohesin exerts distinct functions at different stages of the transcription cycle: it promotes Pol II promoter recruitment while restraining release from pausing (Supplementary Fig. 11). Together, these activities help explain the long-standing paradox that cohesin depletion has only a limited impact on steady-state gene expression, despite severely disrupting 3D genome architecture. Upon cohesin loss, reduced Pol II recruitment and enhanced pause release have compensatory effects, contributing to minimal changes in expression. However, this balance is disrupted upon external stimulation, leading to a marked reduction in transcriptional induction. Our findings indicate that cohesin regulates Pol II recruitment and pausing through distinct mechanisms. To support Pol II loading onto promoters, cohesin facilitates promoter-enhancer communication, as previously proposed, thereby maintaining promoter H3K27ac. In contrast, it transiently engages the transcriptional machinery to ensure an appropriate pausing duration. Sufficient pausing is required for the proper formation of the elongation complex to prevent premature transcription termination. We therefore regard this direct engagement as a quality-control-like step that coordinates elongation factor recruitment and complete transcription.

We found that cohesin ensures proper Pol II dwell time during promoter-proximal pausing. To elucidate the underlying mechanism, our biochemical experiments identified a transcriptional complex at the pausing-to-release transition that includes cohesin, its loader, and Pol II. Notably, a corresponding cohesin- and loader-associated Pol II state was also observed in vivo. These findings suggest that cohesin stabilizes pausing by contributing to the formation of this intermediate complex. In the absence of cohesin, Pol II likely bypasses this step and prematurely enters elongation. Although the precise Pol II pausing position in vitro has not been determined, we concluded that cohesin engages with the transcriptional machinery specifically between pausing and pause release, based on its CDK9 dependency and the co-occurrence of both pausing- and release-associated factors. This interpretation is further supported by two independent approaches: (i) ChIP-seq revealed that cohesin and its loader co-occupy promoters with Pol II, and that their occupancy is sensitive to transcriptional inhibition; and (ii) co-immunoprecipitation demonstrated that CDK9 activity is required for the physical association between the cohesin loader and Pol II. Together, these results suggest that CDK9 modulates the dynamic association between cohesin, its loader, and Pol II across the transcription cycle. Moreover, Inhibition of cohesin’s ATPase activity made this complex detectable, suggesting that it normally disassembles upon cohesin-mediated ATP hydrolysis. We note that ADP–AlFx is a general transition-state analog and does not selectively inhibit the cohesin ATPase. However, the observed effect is cohesin-dependent rather than a consequence of nonspecific ATPase inhibition, as cohesin depletion abolished this complex both in vitro and in vivo. Importantly, while our findings suggest that promoter-associated cohesin contributes to pausing regulation, it remains unclear whether these cohesin molecules also participate in promoter–enhancer loops. Therefore, we cannot exclude the possibility that P–E loops themselves influence pausing, a question that remains to be elucidated.

Proper assembly of elongation factors has been proposed to require sufficient pausing duration^22^. Accordingly, we observed incomplete formation of the elongation complex upon cohesin depletion, potentially due to insufficient pause time. Given that elongation factors facilitate processive and stable transcription by interacting with elongating Pol II^11^, we speculate that cohesin enhances transcription processivity and prevents premature transcription termination by ensuring proper transition from pausing to productive elongation. Consistently, our RNA-seq data revealed enhanced XRN2-mediated RNA surveillance upon cohesin depletion.

Acute cohesin depletion redistributes H3K27ac by reducing it at promoters and increasing it at enhancers, and simultaneously weakens chromatin contacts between these regions. These observations suggest that cohesin mediates long-range promoter–enhancer interactions, which help sustain promoter H3K27ac through enhancer activity (Fig. 6e). Furthermore, these interacting regions span long genomic distances, consistent with previous studies showing that cohesin facilitates the activity of distal enhancers.^32^ A potential mechanism underlying H3K27ac redistribution involves the histone acetyltransferases p300/CBP, which localize at enhancers and deposit H3K27ac not only at enhancers but also at nearby promoters brought into proximity via promoter–enhancer interactions^29,33^. We speculate that cohesin loss reduces this spatial proximity, thereby limiting p300/CBP-mediated acetylation to enhancers. The TFIID complex, a key factor required for Pol II promoter recruitment, also decreased at promoters upon cohesin loss. As inhibition of H3K27ac has been shown to diminish TFIID binding^29^, the reduced H3K27ac likely contributes to the reduction in TFIID occupancy. Together, these findings suggest that cohesin maintains an active chromatin environment and supports efficient Pol II recruitment by sustaining regulatory communication between promoters and enhancers.

Our findings shed light on the molecular basis of CdLS, a developmental disorder caused by cohesin dysfunction. CdLS shares clinical features and a dysregulated transcriptome with CHOPS syndrome, in which SEC component AFF4 is hypermorphic^8,34^. Our results suggest that cohesin stabilizes promoter-proximal pausing and restrains pause release. Thus, aberrantly enhanced pause release may represent a common molecular pathology of CdLS and CHOPS syndrome, as SEC triggers this transition. Consistent with this idea, genome-wide increased AFF4 and decreased NELF at promoters were observed in patient cells from both disorders^34^. This subtle yet widespread dysregulation may become particularly evident in response to external or environmental stimuli during development, as suggested by our findings. Alternatively, an immature elongation complex and destabilized productive elongation may also contribute to disease pathogenesis. Mutations in cohesin factors are also reported in cancers, sometimes co-occurring with additional mutations in other genes.^35,36^ Furthermore, dysregulated gene expression caused by these mutations is proposed to contribute to tumorigenesis, although the underlying mechanism remains unclear. Our findings may oder mechanistic insight into how cohesin dysfunction contributes to transcriptional dysregulation and cancer development.

## Methods

### Cell culture

All HCT116-derived cell lines were cultured in McCoy’s 5A medium supplemented with 10% fetal bovine serum (FBS) and penicillin–streptomycin–L-glutamine (Wako) at 37 °C in a humidified atmosphere containing 5% CO₂.

HCT-116-RAD21-mAID-mClover cells were described previously^12^. RAD21 was depleted by treatment with 0.5 mM indole-3-acetic acid (IAA; Sigma) for 3 h. For CDK9 inhibition, 100 μM 5,6-Dichlorobenzimidazole 1-β-D-Ribofuranoside (DRB; TCI) was added concurrently with IAA for 3 hours, or 1 μM flavopiridol (Santa Cruz) was added during the final hour of IAA treatment. For CDK7 inhibition, 1 μM THZ1 2HCl (Selleck) was added during the final hour of IAA treatment.

HCT116 RAD21-mAID-Clover CMV-OsTIR1(F74G) cells were described previously^37^. RAD21 was depleted by treatment with 1 μM 5-Ph-IAA (Merck Sigma Aldrich) for 2 h.

HCT116 CMV-OsTIR1 SMC1A-mAID and HCT116 Tet-OsTIR1 Stag-3FLAG-NIPBL cells were generated as previously described^12,38^. Briefly, HCT116 CMV-OsTIR1 cells were co-transfected with SMC1A CRISPR and SMC1A-mAID-Hygro donor vectors, and HCT116 Tet-OsTIR1 cells with NIPBL-N CRISPR and Hygro-P2A-Stag-3FLAG-NIPBL donor vectors. After selection with 100 μg/mL HygroGold (Invivogen), single colonies were isolated, and successful knock-in was confirmed by genomic PCR and western blotting. SMC1 was depleted by treatment with 0.5 mM IAA for 3 h in HCT116 CMV-OsTIR1 SMC1A-mAID cells.

HCT116 MAC-NIPBL cells were generated as previously described^12,38^. Briefly, in HCT116 CMV-OsTIR1(F74G) cells, Hygro-P2A-mAID Clover was inserted at the N-terminal coding region of NIPBL. After selection with 100 μg/mL HygroGold (Invivogen), single colonies were isolated, and successful knock-in was confirmed by genomic PCR and western blotting.

### siRNA-mediated knockdown

RNA interference was performed with Lipofectamine RNAiMax (Invitrogen) using siRNA duplexes at a final RNA concentration of 50 nM, according to the manufacturer’s instructions. Cells were collected two days after transfection. The siRNA sequences (5′→3′) were: WAPL, UCUCCUGCUCGUUAGAAGUAAGGG; XRN2, UCGCUAUUACAUAGCUGAUCGUUUA.

### Cell cycle profile analysis

Cells were fixed in 70% ethanol, washed with PBS, treated with RNase A (50 µg/mL), and stained with propidium iodide (50 µg/mL). DNA content was measured on BD Accuri C6 plus flow cytometer.

### Spike-in ChIP-seq and MNase ChIP-seq

ChIP-seq was performed as described previously^39^. MNase ChIP-seq followed the same procedure, except that chromatin was fragmented by sequential micrococcal nuclease (MNase) digestion and sonication. In both assays, mouse C2C12 cells were added as a spike-in control. Briefly, HCT-116 and C2C12 cells were crosslinked with 1% formaldehyde for 10 min, quenched with 125 mM glycine for 5 min, and stored at –80°C. For standard ChIP-seq, chromatin fractionation was performed by sequential resuspension in LB1 Buder [50 mM HEPES-KOH (pH 7.5), 140 mM NaCl, 1 mM EDTA, 10% glycerol, 0.5% NP-40, 0.25% Triton X-100, 1 mM PMSF, 10 mM DTT, 1x cOmplete EDTA-free (Roche)], LB2 Buder [20 mM Tris-HCl (pH 7.5), 200 mM NaCl, 1 mM EDTA, 1 mM PMSF, 1x cOmplete EDTA-free], and LB3 Buder [20 mM Tris-HCl (pH 7.5), 150 mM NaCl, 1 mM EDTA, 1% Triton X-100, 0.1% Na-Deoxycholate, 0.1% SDS, 1x cOmplete EDTA-free]. Chromatin was sonicated on Sonifier 250D (BRANSON). For MNase ChIP-seq, cells were resuspended in MNase Buder [50 mM NaCl, 10 mM Tris-HCl (pH 7.5), 5 mM MgCl_2_, 1 mM CaCl_2_, 0.2% NP-40, 1x cOmplete EDTA-free], digested with MNase (New England BioLabs), and sonicated. HCT-116 and C2C12 chromatin were combined at a 4:1 ratio and immunoprecipitated with antibodies conjugated to Dynabeads Protein A/G (Invitrogen) overnight at 4 °C. DNA was purified using the QIAquick PCR purification kit (QIAGEN), and libraries were prepared with NEBNext DNA Library Prep Master Mix (New England Biolabs) and sequenced on HiSeq 2500 or NextSeq 2000 (Illumina).

The following antibodies were used: Rpb1 NTD (D8L4Y) antibody (Cell Signaling Technology, #14958), Rabbit polyclonal anti-RAD21 (Sakata et al.), Mouse monoclonal anti-MAU2 (Sakata et al.)^34^, Purified Rabbit Anti-GFP antibody (Torrey Pines Biolabs, TP401), Anti-Phospho RNA Polymerase II CTD (Ser2) (Monoclonal research institute Inc., MABI0602), anti-acetyl Histone H3 (Lys27) antibody (MBL International, MABI0309), anti-trimethyl Histone H3 (Lys4) antibody (MBL International, MABI0304), anti-monomethyl Histone H3 (Lys4) antibody (MBL International, MABI0302), Rabbit anti-MCEF Antibody (Bethyl, A302-539A), TH1L (D5G6W) Rabbit mAb (Cell Signaling Technology, #12265), Anti-Cdk9 antibody [EPR22956-37] - ChIP Grade (Abcam, #ab239364), Rabbit anti-TAF1 Antibody, Adinity Purified (Bethyl, #A303-505A), Anti-TATA binding protein TBP antibody [mAbcam51841] (Abcam, #ab300656).

### Quantitative PCR

ChIP DNA was quantified using 7500 and StepOnePlus real-time PCR systems (Applied Biosystems) with KAPA SYBR Fast qPCR kit (KAPA Biosystems). The following primers were used (5′→3′): Rad21_1 F: GGCCTAGAAAACACGTCTACAG, R: TCCACCAAGGGACAAACAGAG; Rad21_2 F: TGCGGGTACACTTTGAGAACC, R: GTGCCACCTAGTGGTATTTCCT; Rad21_3 F: CTTTGCCCCTTTGTGGCTCA, R: ATCCTGACACTGCTGGGAATG; negative control F: ATGAGTCTGGGATTTGGCTGG, R: TCTCAGGGGAAGGAGAGAGTC.

### RNA-seq

Total RNA was extracted from HCT-116 cells with TRIzol reagent (Invitrogen) and purified with NucleoSpin RNA II kit (Macherey-Nagel), following the manufacturer’s instructions. rRNA was depleted with NEBNext rRNA Depletion Kit (New England BioLabs). RNA-seq libraries were prepared using NEBNext Ultra II Directional RNA Library Prep Kit for Illumina and NEBNext Multiplex Oligos for Illumina (New England BioLabs). The libraries were sequenced on Nextseq 2000 (Illumina).

### TT-seq

TT-seq was performed as described previously with minor modifications^40^. Drosophila S2 cells were used as a spike-in control. Nascent transcripts were labelled by incubating HCT-116 cells and S2 cells with 1 mM 4-thiouridine (Cayman Chemical) for 10 min and 20 min, respectively. Total RNA was extracted with TRIzol reagent (Invitrogen) and purified using Direct-zol RNA Miniprep Plus (ZYMO RESEARCH). For each library, 100 μg of HCT-116 RNA was mixed with 1 μg S2 RNA and sheared on M220 Focused-ultrasonicator (Covaris). Biotinylation of 4sU-labeled RNA was carried out with MTSEA-biotin-XX (20 mg/mL), followed by phenol–chloroform extraction and purification using the mMACS Streptavidin Kit (Miltenyi Biotec) and Direct-zol RNA Miniprep. TT-seq libraries were generated with the NEBNext Ultra II Directional RNA Library Prep Kit and Multiplex Oligos (New England Biolabs) and sequenced on a NextSeq 2000 (Illumina).

### EU-seq

Drosophila S2 cells were used as a spike-in control. Nascent transcripts were labelled with 0.5 mM 5-ethynyl uridine (Thermo Fisher) for 20 min (HCT-116) or 0.2 mM for 2 hours (S2). Total RNA was extracted with TRIzol reagent (Invitrogen) and purified using NucleoSpin RNA II kit (Macherey-Nagel). rRNA depletion of HCT-116 RNA was performed with the NEBNext rRNA Depletion Kit (New England Biolabs), and S2 mRNA was purified with the Dynabeads mRNA Purification Kit (Invitrogen). For each library, 200 ng of HCT-116 RNA was mixed with 20 ng of S2 mRNA, and EU-labelled RNA was isolated with Click-iT Nascent RNA Capture Kit (Thermo Fisher). cDNA was synthesized with SuperScript VILO cDNA synthesis kit (Thermo Fisher). Second-strand cDNA synthesis and library preparation were performed with NEBNext Ultra II Directional RNA Library Prep Kit for Illumina and NEBNext Multiplex Oligos for Illumina (New England BioLabs). The library was sequenced on Nextseq 2000 (Illumina).

### mNET-seq

mNET-seq was performed as described previously with minor modifications^41^. Briefly, chromatin fractionation from 1 × 10^8^ HCT-116 cells were performed using HLB+N buder [10 mM Tris-HCl (pH 7.5), 10 mM NaCl, 2.5 mM MgCl_2_, 0.5% NP-40, 1x cOmplete EDTA-free (Roche), 1x PhosSTOP (Roche)], HLB+NS buder [10 mM Tris-HCl (pH 7.5), 10 mM NaCl, 2.5 mM MgCl_2_, 0.5% NP-40, 1x cOmplete EDTA-free, 1x PhosSTOP, 10% sucrose], NUM1 lysis buder [20 mM Tris-HCl (pH 7.9), 75 mM NaCl, 0.5 mM EDTA (pH 8.0), 50% glycerol, 1x cOmplete EDTA-free, 1x PhosSTOP], and NUM2 lysis buder [20 mM HEPES (pH 7.5), 300 mM NaCl, 0.2 mM EDTA (pH 8.0), 7.5 mM MgCl_2_, 1% NP-40, 1M urea, 1x cOmplete EDTA-free, 1x PhosSTOP]. Chromatin was digested with MNase in MNase reaction buder [1x MNase buder, 1x BSA, 400 gel units/μL MNase (New England BioLabs)]. Pol II complexes were immunoprecipitated using Pol II CTD antibody (mAb) (Monoclonal research institute Inc.) and Dynabeads Protein G (Invitrogen). After washes, bead-bound RNA was phosphorylated with T4 PNK (NEB), extracted with TRIzol, and purified using Direct-zol RNA Miniprep (Zymo Research). Drosophila S2 mRNA was prepared as a spike-in control by extraction, poly(A) selection, fragmentation, and phosphorylation, and 2.5 ng was added to each mNET-seq sample. Libraries were constructed using the NEBNext Small RNA Library Prep Set (NEB) with size selection on AMPure XP beads (Beckman Coulter) and sequenced on a NextSeq 550 (Illumina).

### Micro-C

Micro-C was performed as described previously^34^. Briefly, 5×10^6^ cells were crosslinked with 1% formaldehyde for 10 min and quenched with 125 mM glycine for 5 min. After PBS washing, cells were further crosslinked with 3 mM DSG (Thermo Fisher) for 40 min, quenched with 400 mM glycine for 5 min, and stored at –80°C. Cells were resuspended in Micro-C Buder #1 [50 mM NaCl, 10 mM Tris-HCl (pH 7.5), 5 mM MgCl_2_, 1 mM CaCl2, 0.2% NP-40, 1x cOmplete EDTA-free (Roche)] and digested with MNase. DNA ends were repaired, filled in with biotinylated nucleotides, and ligated with T4 ligase (NEB). Unligated DNA ends were digested with Exonuclease III (NEB). Following proteinase K treatment and reverse cross-linking, DNA was purified with QIAquick PCR Purification Kit (QIAGEN) and size-selected with AMPure XP beads and subjected to biotin pull-down with Dynabeads MyOne Streptavidin C1 (Invitrogen). End repair, adaptor ligation, and PCR amplification were carried out on beads using the NEBNext Ultra II DNA Library Prep Kit and Multiplex Oligos (NEB) according to the manufacturer’s instructions. Libraries were sequenced on a NovaSeq 6000 (Illumina).

### Digitonin permeabilization and CUT&Tag

We used HCT116 CMV-OsTIR1 (F74G) RAD21-mAID-mClover cells for RAD21 (with anti-GFP antibody) and RPB1 CUT&Tag and used HCT116 CMV-OsTIR1 (F74G) mAID-mClover-NIPBL cells for NIPBL CUT&Tag (with anti-GFP antibody).

Cells were detached from a culture dish by Accutase (Thermo Fisher Scientific) and incubated on ice for 15 min in dig-wash buder [20 mM HEPES-KOH (pH 7.5), 150 mM NaCl, 1 mM EDTA, 0.05% digitonin, 0.5 mM spermidine, 1 mM DTT, 1x cOmplete EDTA-free (Roche)] for permeabilization. The cells were washed twice with Dig-wash buder and resuspended in reaction buder [20 mM HEPES-KOH (pH 7.5), 75 mM NaCl, 5 mM MgCl2, 0.05% digitonin, 10% glycerol, 1 mM DTT, 1x cOmplete EDTA-free] and incubated at 25°C for 10 min with or without 1 mM ATP. For AlFx, 1 mM AlCl3 and 5 mM NaF were added after the incubation with 1 mM ATP for 5 min and the cells were further incubated for 5 min.

CUT&Tag was performed as previously described with minor modifications^42,43^. Briefly, cells were crosslinked with 0.3% formaldehyde for 2 min, quenched with 225 mM glycine for 5 min. The crosslinked cells were washed once with cold PBS and resuspended in wash buder [20 mM HEPES-KOH (pH 7.5), 150 mM NaCl, 0.5 mM spermidine, 1× cOmplete EDTA-free (Roche)]. Concanavalin A–conjugated magnetic beads were prepared by incubating Dynabeads MyOne T1 (Invitrogen) with biotinylated concanavalin A (Type IV; Merck Sigma-Aldrich) in bead wash buder [PBS (pH 6.8) containing 0.01% Tween-20]^44^. Cells were immobilized on concanavalin A–coated magnetic beads and incubated in antibody buder [wash buder supplemented with 0.1% Triton X-100, 2 mM EDTA, and 100 μg mL⁻¹ BSA (NEB)] containing the appropriate primary antibody for 2 h at room temperature, followed by incubation with secondary antibody for 1 h at room temperature. Cells were washed once with TX-wash buder [wash buder containing 0.1% Triton X-100] and once with 300-TX-wash buder [TX-wash buder containing 300 mM NaCl], and then incubated with CUTANA™ pAG-Tn5 (EpiCypher). Tagmentation was performed at 37 °C for 1 h in tagmentation buder [300-TX-wash buder supplemented with 10 mM MgCl2 and 5 ng mL⁻¹ CUTANA™ E. coli spike-in DNA (EpiCypher)]. DNA was released in 2x release buder [20 mM HEPES-KOH (pH 7.5), 20 mM EDTA, 2% SDS, and 2.5 μg mL⁻¹ proteinase K (Roche)] and incubated at 55 °C for 2 h. DNA was purified using a QIAquick PCR purification kit (QIAGEN), amplified with indexed primers^45^, size-selected using SPRIselect beads (Beckman Coulter), and sequenced on a NextSeq 1000 (Illumina).

The following antibodies were used: Rabbit Anti-GFP antibody (abcam, #ab290), Rpb1 CTD (4H8) Mouse Monoclonal Antibody (Cell Signaling Technology, #2629), Anti-Rabbit Secondary Antibody for CUTANA™ CUT&Tag Workflows (EpiCypher, #1047), Rabbit Anti-Mouse IgG H&L (abcam, #ab46540).

### Co-immunoprecipitation

Co-immunoprecipitation was performed using the nuclear fraction. Cells were washed in buder A [20 mM Tris-HCl (pH 7.5), 10 mM NaCl, 1.5 mM MgCl_2_, 10% glycerol, 10% sucrose, 1x cOmplete EDTA-free (Roche), 1x PhosSTOP (Roche), 0.2% Triton X-100] twice and stored at −80°C. The frozen pellet was lysed in lysis Benzonase buder [20 mM Tris-HCl (pH 7.5), 150 mM NaCl, 0.05% NP-40, 10% glycerol, 2.5 mM MgCl_2_, 1x cOmplete EDTA-free, 1x PhosSTOP, Benzonase (Millipore)], and centrifuged. The pellet was further resuspended in another lysis buder [20 mM Tris-HCl (pH 7.5), 250 mM NaCl, 0.1% NP-40, 10% glycerol, 1x cOmplete EDTA-free, 1x PhosSTOP]. Supernatants from both lysis steps were pooled and incubated with antibody-conjugated beads for 3 hours at 4°C. Beads were washed five times with buder [20 mM Tris-HCl (pH 7.4), 200 mM NaCl, 5% glycerol, 1 mM EDTA, 0.1% NP-40], and bound proteins were eluted in SDS sample loading buder and boiled for 5 min at 95°C.

The following antibodies were used for co-immunoprecipitation: Anti-RNA Polymerase II CTD mAb (Fujifilm, #384-09271), Anti-Phospho RNA Polymerase II CTD (Ser2) mAb (Fujifilm, #381-09281), Anti DYKDDDDK tag Antibody Magnetic Beads (Fuijifilm, #017-25151), Nipbl (Short form) Rat Monoclonal Antibody [Clone ID: KT55] (ORIGENE, #AM20797PU-N).

### *In vitro* transcription assay

Nuclear extracts from HeLa S3 cells were prepared as described by Dignam et al^46^. Cohesin was immunodepleted twice by incubating nuclear extract with protein A magnetic beads conjugated to anti-SMC1 and anti-SMC3 antibodies. The Mediator complex was immunodepleted twice using protein G magnetic beads conjugated to anti-MED4, anti-MED15, and anti-MED25 antibodies. A plasmid (p5G-sMLT) was constructed by inserting five tandem GAL4 binding sites, an 81-bp minimal promoter region containing a TATA box, and a 245-bp *slc16a1* first intron fragment into the pUC19 multicloning site. DNA templates were amplified using a biotinylated primer 50 bp upstream of the GAL4 sites and a reverse primer 419 bp downstream of the TATA box, and conjugated to Dynabeads M-280 Streptavidin (Invitrogen) in binding buder [10 mM Tris-HCl (pH 7.5), 1 M NaCl, 0.05% Tween-20]. After washing, templates were incubated with GAL4-VP16 in activator binding buder [12.5 mM HEPES-KOH (pH 7.4), 62.5 mM KCl, 7.5 mM MgCl₂, 12.5% glycerol, 0.025% NP-40, 100 μg/mL BSA, 0.625 mM DTT]. After washing using activator binding buder without BSA, GAL4-VP16-bound templates were incubated with nuclear extracts in reaction buder [12.5 mM HEPES-KOH (pH 7.4), 62.5 mM KCl, 7.5 mM MgCl₂, 12.5% glycerol, 0.025% NP-40, 100 μg/mL BSA, 0.625 mM DTT, 2.5 mM NU-7441, 0.5x cOmplete EDTA-free] at 23°C. AlFx and 100 μM NTPs or ADP, prepared by mixing 1 mM AlCl₃ and 5 mM NaF, were added to the reaction solution. The timing of addition is described in the corresponding figure legend. After the reaction, the beads were washed three times with activator binding buder without BSA and eluted in SDS sample loading buder.

To detect RNA production, beads conjugated with biotinylated DNA templates, either pre-bound or not with GAL4-VP16, were incubated with HeLa nuclear extract in the presence of an RNase inhibitor (RNasin® Ribonuclease Inhibitor, Promega) for 20 min at room temperature. After incubation for 30 min at 30 °C, reactions were terminated by adding an equal volume of stop solution (0.5% SDS, 10 mM EDTA).

Supernatants were collected by centrifugation at 10,000 × g for 1 min. RNA was purified using the Direct-zol™ RNA Microprep kit according to the manufacturer’s instructions. Residual template DNA was removed by treatment with ezDNase (Invitrogen) for 2 min at 37 °C. Reverse transcription was performed using SuperScript™ III Reverse Transcriptase (Invitrogen) and a gene-specific primer (5′→3′) (GGATAACAATTTTCACACAGG), which anneals 5 bp upstream of the 3’ end of the transcript. Quantitative PCR was performed using 7500 and StepOnePlus real-time PCR systems with KAPA SYBR FAST qPCR Master Mix. The following primers were used (5′→3′): Site 1 F: GCCAGTGAATTCGAGCTCGGTA, R: GAAGAACTAGTTGCGGCCGCTGCTAGC; Site 2 F: CGGGGTTGGGAGACTTTGTC, R: GCCTCAAGGTCTCCTTCACC; Site 3 F: CTCTCTTAGCCGAGCCGC, R: TTCCTCGAGTGGAAGGGAC. Relative RNA levels were quantified using standard curves generated from the full-length template DNA.

### Purification of recombinant proteins

NIPBL mutants N1 (1–700) and N2 (701–1000) were N-terminally FLAG-tagged and subcloned into the pFASTBac HTB vector by GenScript. NIPBL mutant C (1121–2804) was C-terminally FLAG-tagged and subcloned into the pAcHLT-B vector, also by GenScript. Wild-type MAU2 was subcloned into the pAcSG2 vector by GenScript. Recombinant baculoviruses were generated using either the flashBAC Baculovirus Expression System (Mirus) or the Bac-to-Bac Baculovirus Expression System (Gibco), according to the manufacturer’s protocols. NIPBL fragments (N2 and C), as well as N1 co-expressed with MAU2, were expressed in Sf9 cells following infection with recombinant baculoviruses. Cells were lysed using a Dounce homogenizer in PB500 buder [25 mM NaH₂PO₄/Na₂HPO₄ (pH 7.5), 500 mM NaCl, 5% glycerol] supplemented with 10 mM imidazole, 0.05% Tween-20, 0.1% NP-40, 1 mM PMSF, 3 mM 2-mercaptoethanol, and 1x cOmplete EDTA-free (Roche). Lysates were clarified by centrifugation at 35,000 × g for 1 hour and incubated with Ni-NTA Agasore (Qiagen) for 2 hours at 4 °C. The resin was washed with PB500 supplemented with 20 mM imidazole, and bound proteins were eluted with PB150 (same composition as PB500 but with 150 mM NaCl) supplemented with 300 mM imidazole. Eluted proteins were further purified by incubation with ANTI-FLAG M2 Adinity Gel (Sigma) for 2 hours at 4 °C, washed with PB150, and eluted with PB150 containing 0.5 mg/ml 3×FLAG peptide. Final protein preparations were concentrated and buder-exchanged into 20 mM Tris-HCl (pH 7.5), 100 mM NaCl, and 20% glycerol using Amicon Ultra Centrifugal Filters, 50kDa MWCO (Millipore).

### SDS-PAGE, CBB staining, and western blotting

Protein samples were resolved by SDS–PAGE using either 4–15% Mini-PROTEAN® TGX™ Precast Protein Gels (Bio-Rad) or Perfect NT precast gels (DRC). For Coomassie Brilliant Blue (CBB) staining, gels were stained with one-step CBB stain solution (BIO CRAFT). For western blotting, proteins were transferred to Amersham Protran 0.2 NC nitrocellulose Western blotting membranes (GE Healthcare). The membranes were blocked with 3% skim milk in PBS-T and incubated with primary antibodies for several hours or overnight. After washing with PBS-T, the membranes were incubated with HRP-conjugated secondary antibodies for 1 hour. Signals were detected using SuperSignal West Pico PLUS Chemiluminescent Substrate (Thermo Scientific). When required, membranes were stripped using Stripping Solution (Wako) and reprobed.

### Spike-in ChIP-seq and MNase ChIP-seq alignment and data analysis

Reads were aligned to human (hg38) and mouse (mm10) genomes with Bowtie (version 1.2.3)^47^. Normalization factors were calculated from the ratio of mouse spike-in to human reads in ChIP (IP) and input (WCE) samples, as described previously^34^. Redundant reads were removed, and read numbers were normalized to a spike-in control with parse2wig (version 3.7.2)^48^. Peak calling was performed with drompa_peakcall (version 3.7.2) in “PC_SHARP” mode^48^. Visualization was carried out with drompa+ PC_SHARP (version 1.17.2) and deepTools(version 3.5.1)^49,50^.

### RNA-seq alignment and data analysis

Reads were aligned to the human genome (hg38) using STAR (version 2.7.10a)^51^. Transcript abundance was quantified using RSEM (version 1.3.3)^52^. Differential expression analysis was performed with edgeR (version 3.40.2; FDR < 0.05)^53^.

### TT-seq alignment and data analysis

Reads were aligned to human (hg38) and Drosophila (dm6) genomes with STAR (version 2.7.1a). Mapped reads were divided into sense and anti-sense reads using samtools (version 1.15.1)^54^. Scale factors were calculated from a Drosophila gene-level count matrix generated with htseq-count (version 1.99.2) and normalized with DESeq2’s estimateSizeFactors function^55,56^. A read count table was generated for genes, with TT-seq reads counted using drompa3 (version 3.7.2)^48^, and differential expression analysis was performed with edgeR (version 4.2.2) with the likelihood ratio test (FDR < 0.05)^53^.

### EU-seq alignment and data analysis

Low-quality reads were removed with Trimmomatic (version 0.39)^57^. Reads were aligned to the Drosophila genome (dm6) using Bowtie2 (version 2.4.5) in “--very-sensitive-” mode, and the unmapped reads were aligned to the human genome (hg38)^58^. Scale factors were calculated from a Drosophila gene-level count matrix generated with htseq-count (version 1.99.2) and normalized with DESeq2’s estimateSizeFactors function. Redundant reads were removed, and the remaining reads were normalized to a spike-in control with parse2wig (version 3.7.2). Differential expression analysis was performed with edgeR (version 4.2.2) with the likelihood ratio test (FDR < 0.05)^53^.

### mNET-seq alignment and data analysis

Reads were aligned to human (hg38) and Drosophila (dm6) genomes with STAR (version 2.7.1a). Post-processing steps, including read deduplication, RNA 3’ end identification, removal of PCR internal priming, and exclusion of reads associated with co-transcriptional splicing, were performed as described by Prudêncio et al^59^. Scale factors were calculated from a Drosophila gene-level count matrix generated with htseq-count (version 1.99.2) and normalized with DESeq2’s estimateSizeFactors function. Mapped reads were divided into sense and anti-sense reads using samtools^54^ (version 1.15.1).

### Micro-C alignment and data analysis

Reads were aligned to the human genome (hg38) with BWA-MEM (version 0.75)^60^. Ligation-junction detection, read sorting, PCR-duplicate removal, and pair-file generation were performed using Pairtools (version 0.3.0)^61^. Matrix files were generated from the pairs files using Juicer (version 1.22.01) at 5 kb, 2 kb, and 1 kb resolutions. Loops were identified using Juicer Tools HiCCUPS and Mustache (version 1.2.0), and loops detected by both methods were combined to generate a set of Micro-C loops ^62,63^. TADs were identified using Juicer Tools Arrowhead at a 25 kb resolution. Loop intensity was quantified using Chromosight^64^ (version 1.6.1). For visualization, contact maps were generated using HiCExplorer (version 3.7.2), and aggregate plots were created using coolpup.py (version 1.0.0)^65,66^.

### CUT&Tag alignment and data analysis

Reads were aligned to human (hg38) and E. coli genomes with Bowtie2 (version 2.5.3) using Churros (version 1.5.0)^67^. Normalization factors were calculated from the ratio of E. coli spike-in to human reads. Redundant reads were removed, and read counts were normalized by the normalization factors with parse2wig+ (version 1.20.0). Visualization was carried out with drompa+ PC_SHARP (version 1.17.2) and deepTools(version 3.5.1).

### Identification of +1 nucleosome and pausing site

Genomic locations of nucleosomes with H3K4me3 modifications were identified from H3K4me3 MNase ChIP-seq data with DANPOS3 (version 2.2.2)^68^. Nucleosomes located between TSS +10 bp and TSS + 300 bp with an occupancy score > 1, a P-value for occupancy < −1, a fuzziness score < 100, and a P-value for fuzziness < −2 were annotated as +1 nucleosomes.

Pausing sites were defined as the genomic position between the TSS and the +1 nucleosome dyad with the highest mNET-seq signal. mNET-seq signal intensity was quantified in 1-bp bins and smoothed using a moving average with a window size of 11. The intensity at the pausing site was required to exceed 1.2 times the median intensity of the surrounding regions.

### Selection of genes for analysis

We selected 3,597 genes that met all of the following criteria: TT-seq TPM ≥ 1, gene length ≥ 5 kb, and >1 kb from neighboring genes, and the presence of a +1 nucleosome and a defined pausing site.

### Calculation of the pausing-duration index, Pol II ChIP-seq pausing index, the half-time of paused Pol II, velocity index, and the slope of RNA-seq intronic reads

The pausing-duration index was calculated as the mNET-seq/TT-seq signal ratio around pausing sites for each gene. TT-seq and mNET-seq signals within ± 25 bp of pausing sites were quantified using multiBigwigSummary from deepTools.

Pol II ChIP-seq pausing index was calculated as the ratio of RPB1 occupancy at promoters to that at gene bodies. RPB1 occupancy at promoters (TSS −500 bp to 500 bp) and at gene bodies (TSS +2 kb to +3 kb) was quantified using multiBigwigSummary from deepTools.

The half-time of paused Pol II was estimated by applying a single-exponential decay model to the reduction in RPB1 occupancy at promoters over the time-course experiment following triptolide treatment. RPB1 ChIP-seq was performed 0, 5, 10, and 20 min after triptolide treatment, and RPB1 occupancy at promoters (TSS – 0.5 kb to + 0.5 kb) was quantified using multiBigwigSummary from deepTools. Genes exhibiting a continual reduction in RPB1 occupancy throughout the time-course experiment were analyzed using a single-exponential decay model fitted with nls() in R.

The velocity index was defined as the TT-seq/mNET-seq signal ratio along genes. mNET-seq and TT-seq signals were quantified using multiBigwigSummary from deepTools.

RNA-seq read intensity at introns longer than 1 kb was quantified using multiBigwigSummary from deepTools with 100-bp bins. A simple linear regression model was fitted to the quantified intensity of each intron using lm() in R. Introns with a negative slope and adjusted R^2^ ≥ 0.5 were analyzed.

### K-means clustering of genes

K-means clustering (k=3) was performed in R on the log₂FC of promoter H3K27ac, pausing-duration, velocity index, and TT-seq gene expression for each gene. All indices were normalized before clustering.

### Identification of promoter and enhancer regions

Twelve chromatin states were defined in control and auxin-treated cells using ChromHMM (version 1.25)^30^ with ChIP-seq datasets for histone modifications (H3K27ac, H3K4me1, H3K4me3, H3K36me3, H3K79me2, H3K4me2, H2Aub, H2Bub, H3K27me3, H3K9me3), chromatin-associated factors (CTCF, BRD4, MED1), and TT-seq signal, all obtained from both conditions. Chromatin segments enriched for H3K27ac, H3K4me3, BRD4, and MED1 were annotated as active promoters, while those enriched for H3K27ac, H3K4me1, BRD4, and MED1 were classified as active enhancers.

### Identification of up- or down-regulated promoters and enhancers

Active promoters and enhancers defined by ChromHMM were merged, and H3K27ac signal intensity was compared between control and auxin-treated cells using a binomial test with drompa_draw CI. FDRs were computed from p-values using p.adjust in R, and peaks with FDR < 0.05 were classified as increased or decreased based on signal changes.

### Evaluation of reproducibility

Reproducibility between biological replicates was systematically assessed for TT-seq, RNA-seq, EU-seq, ChIP-seq, and mNET-seq. Reproducibility was evaluated by calculating Spearman correlations of genome-wide read coverage in 10-kb bins after excluding bins containing zero counts. For Micro-C, replicate reproducibility was quantified using QuASAR-Rep scores as implemented in 3DChromatin_ReplicateQC at 100-kb resolution^69^. All assays showed high reproducibility across conditions, with replicate correlations and QuASAR-Rep scores reported in Supplementary Tables 1and 2.

### Simulation of Pol II State Transitions

We modeled transcription dynamics under cohesin depletion using a discrete-time Markov chain, in which Pol II transitions among four states: Free (1), Initiation (2), Paused (3), and Elongation (4). (Supplemental Fig. 10a). Pol II progresses sequentially through Initiation and Paused before Elongation, but can return directly to the Free from either Initiation or Paused. Dwell times in the Free, Initiation, and Paused states were variable and modeled through self-transitions, whereas Elongation had a fixed hold time (*θ₄*) before returning to Free.

We assigned the numbers 1-4 to the states Free, Initiation, Paused, and Elongation, respectively. Letting *p*_ij_ denote the transition probability from state *i* to state *j*,

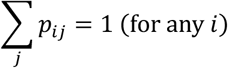

Considering the constraints above, the transition probability matrix *P* is:

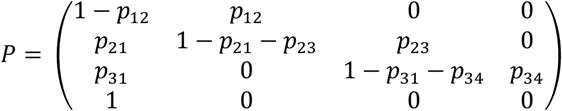

This model has six independent parameters: five transition probabilities (*p*_12_, *p*_21_, *p*_23_, *p*_31_, *p*_34_), and the hold time *θ*_4_ in the Elongation state.

Next, we calibrated the parameters to reflect the actual *in vivo* behavior of Pol II reported previously^31^. One step in the Markov model was assumed to correspond to one second. The mean durations in the Initiation and Paused states were

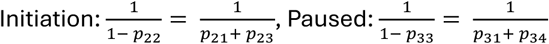

Reported ratios of *p*_21_:*p*_23_, and *p*_31_:*p*_34_ were used to determine *p*_21_, *p*_23_, *p*_31_ and *p*_34_, and *8*_4_ was also set to match empirical measurements. (Supplemental Fig. 10b)

The remaining parameter *p*_12_ was determined to reproduce the observed stationary distribution. Since the transition probability matrix has a left eigenvector with eigenvalue 1, we let this eigenvector be (*α*_1_, *α* _2_, *α* _3_, *α*_4_). The stationary distribution is then

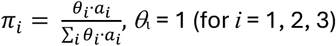

*p*_12_ was chosen to minimize the diderence between the calculated and empirical distribution. All model simulations were conducted using R.

## Data availability

Micro-C, ChIP-seq, TT-seq, mNET-seq, RNA-seq, and EU-seq data were deposited in NCBI Gene Expression Omnibus (GEO) under the following accession numbers: GSE260854 (Micro-C), GSE260852 and GSE300575 (ChIP-seq), GSE300574 (TT-seq), GSE300576 (mNET-seq), GSE300577 (RNA-seq), and GSE260853 (EU-seq). All data are publicly available.

## Acknowledgements

We would like to thank all members of Shirahige Laboratory for discussions. This work was supported by JST CREST (grant number JPMJCR18S5), JSPS KAKENHI (grant numbers JP20H05940, JP20H05933, JP20H05686, and JP25H00969), AMED BINDS (grant number 22ama121020j0001), AMED ASPIRE-A (grant number JP23jf0126003), and the Swedish Research Council (registration number 2022-03478). All grants were awarded to K.S.

## Author Contributions

S.T., M.B., T.Sa., and K.S. designed the study. S.T. performed Micro–C, TT–seq, mNET–seq, and most ChIP–seq experiments, led the genome-wide analyses, integrated the datasets, wrote the original draft, and coordinated the revision. M.B. contributed to ChIP–seq experiments and performed immunoprecipitation and in vitro transcription assays. T.Sa. performed CUT&Tag experiments. A.Y. performed RNA–seq experiments. T.Su. conducted transcription-cycle simulations and contributed to computational analyses. T.N. and M.T.K. provided key resources. K.S. supervised the project and acquired funding. All authors reviewed and edited the manuscript.

## Competing Interests statement

The authors declare no competing interests.

## Notes

### Competing Interest Statement

The authors have declared no competing interest.

### Summary of Updates

New in vivo CUT&Tag under ATP→ADP-AlFx conditions supports the physiological existence of the cohesin-loader-Pol II promoter complex identified in vitro and advances our mechanistic understanding of how cohesin regulates Pol II pausing. Additional ChIP-seq analyses enabled a more precise assessment of the effects of cohesin depletion, revealing that cohesin promotes Pol II promoter recruitment rather than transcription initiation. Reanalysis of published datasets (Rao et al.) from the same cell line independently strengthens our conclusions.

