## Supplemental figures and tables for "Cohesin acts as a transcriptional gatekeeper by restraining pause–release to promote processive elongation"

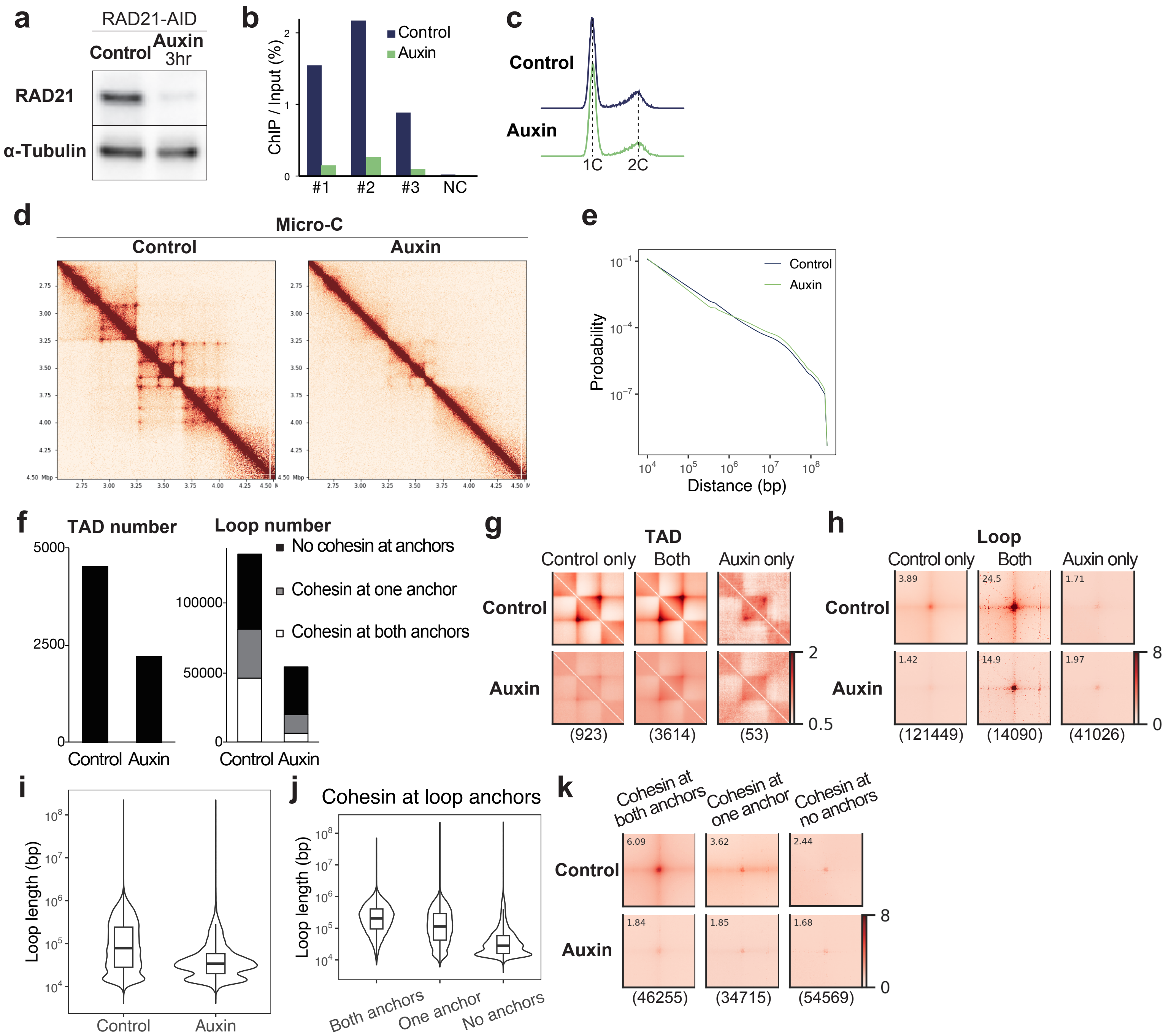


**Supplementary Fig. 1: Cohesin is a major determinant of 3D chromatin structure.**

**a** Western blots of RAD21 and α-tubulin in whole-cell extracts. HCT-116 cells expressing AID-tagged RAD21 were treated with 0.5 mM indole-3-acetic acid (auxin) for 3 hours. RAD21 protein levels, normalized to α-tubulin, were reduced to ~6% of control levels upon auxin treatment.

**b** ChIP–qPCR of RAD21 at three RAD21 ChIP–seq peaks and a negative control (NC) site. An anti-GFP antibody was used to detect RAD21 fused at its C-terminus with a mini auxin-inducible degron (mAID) and mClover.

**c** Histograms of DNA content in control and auxin-treated cells. Fixed cells were stained with propidium iodide, and DNA content was measured by flow cytometry after 3-h auxin treatment.

**d** 5-kb resolution Micro-C contact maps in control and auxin-treated cells. Shown is a 2-Mb region on chromosome 18.

**e** Contact probability curves derived from Micro-C in control and auxin-treated cells.

**f** Bar plots showing the number of TADs and loops identified by Micro-C in control and auxin-treated cells. TADs were called at 25-kb resolution. Loops were detected at 1-, 2-, and 5-kb resolutions, merged, and categorized by the presence or absence of RAD21 peaks at loop anchors.

**g** Aggregate plots of TADs detected only in control cells, only in auxin-treated cells, or shared between both conditions. The number of TADs in each category is indicated in parentheses.

**h** Aggregate plots of loops detected only in control cells, only in auxin-treated cells, or shared between both conditions. The number of loops in each category is indicated in parentheses.

**i** Violin plots showing the distribution of loop lengths derived from Micro-C in control and auxin-treated cells.

**j** Violin plots showing the distribution of loop lengths in control cells, categorized by the presence or absence of RAD21 peaks at loop anchors.

**k** Aggregate plots of loops categorized by the presence or absence of RAD21 peaks at loop anchors. The number of loops in each category is indicated in parentheses.

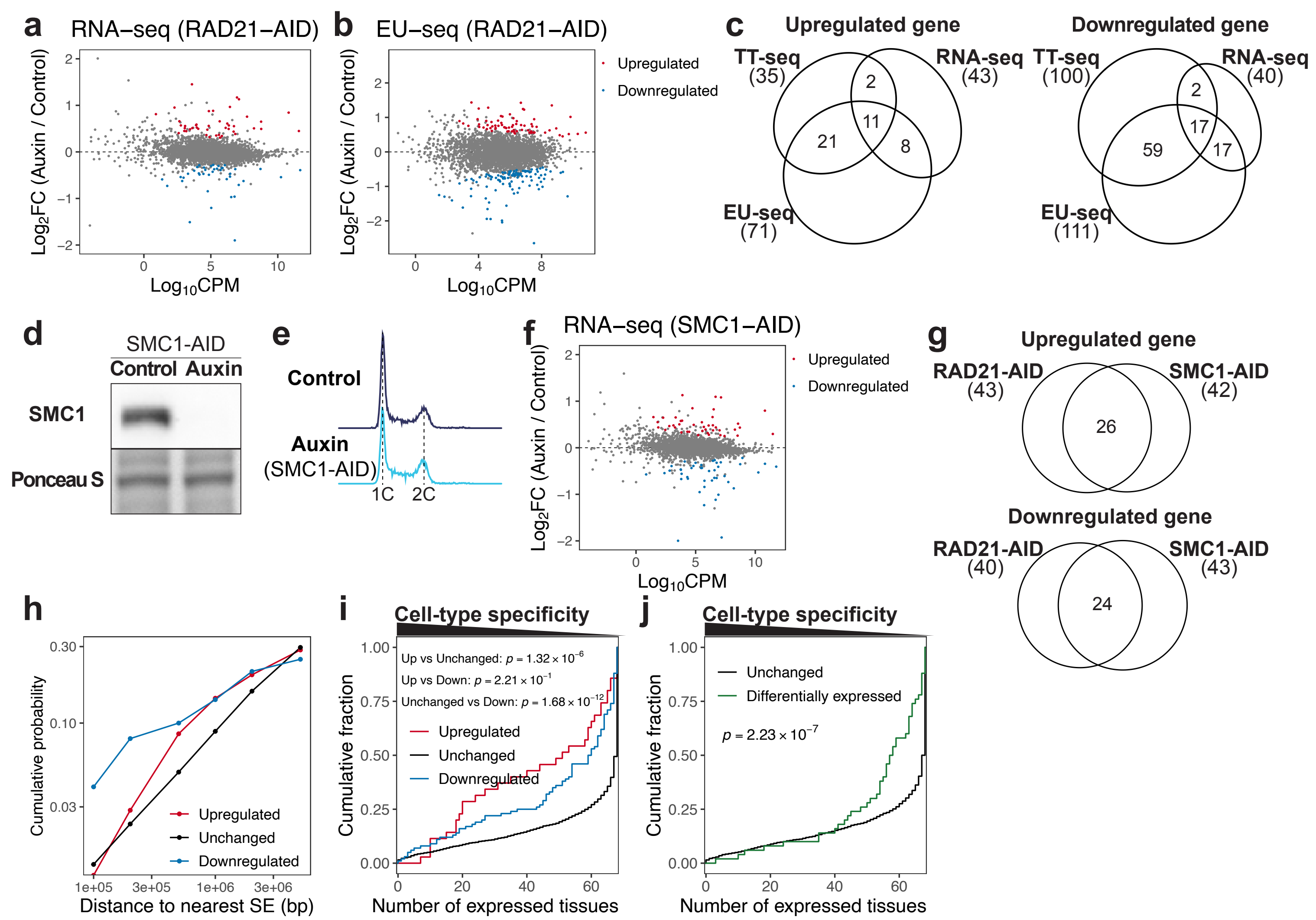


**Supplementary Fig. 2: Cohesin loss has limited effects on steady-state gene expression**

**a** MA plot of RNA-seq showing gene expression changes upon auxin-induced cohesin depletion (3,597 genes). Differentially expressed genes were identified using edgeR (FDR < 0.01) based on three biological replicates.

**b** MA plot of EU-seq showing gene expression changes upon auxin-induced cohesin depletion (3,597 genes). Differentially expressed genes were identified using edgeR (FDR < 0.05) based on two biological replicates.

**c** Venn diagrams showing the overlap among upregulated and downregulated genes identified from TT-seq, RNA-seq, and EU-seq.

**d** Western blots of SMC1 and Ponceau staining in whole-cell extracts. HCT-116 cells expressing AID-tagged SMC1 were treated with 0.5 mM indole-3-acetic acid (auxin) for 3 h. SMC1 protein levels, normalized to Ponceau, decreased to ~1% of control levels upon auxin treatment.

**e** Histograms of DNA content in control and auxin-treated cells expressing SMC1-AID. Fixed cells were stained with propidium iodide, and DNA content was measured by flow cytometry after 3-hour auxin treatment.

**f** MA plot of RNA-seq showing gene expression changes upon auxin-induced cohesin (SMC1) depletion (3,597 genes). Differentially expressed genes were identified using edgeR (FDR < 0.01) based on three biological replicates.

**g** Venn diagrams showing the overlap between upregulated and downregulated genes identified from RNA-seq in RAD21-depleted and SMC1-depleted cells.

**h** Cumulative probability distributions of distances to the nearest super enhancers for genes upregulated, unchanged, and downregulated upon cohesin depletion identified by TT-seq. Statistical significance was calculated using the Kolmogorov–Smirnov test.

**i** Cumulative distributions of cell-type specificity for genes upregulated, unchanged, and downregulated upon cohesin depletion identified by TT-seq. Statistical significance was calculated using the Kolmogorov–Smirnov test.

**j** Cumulative distributions of cell-type specificity for differentially expressed genes shared between RAD21-depleted and SMC1-depleted cells. Statistical significance was calculated using the Kolmogorov–Smirnov test.

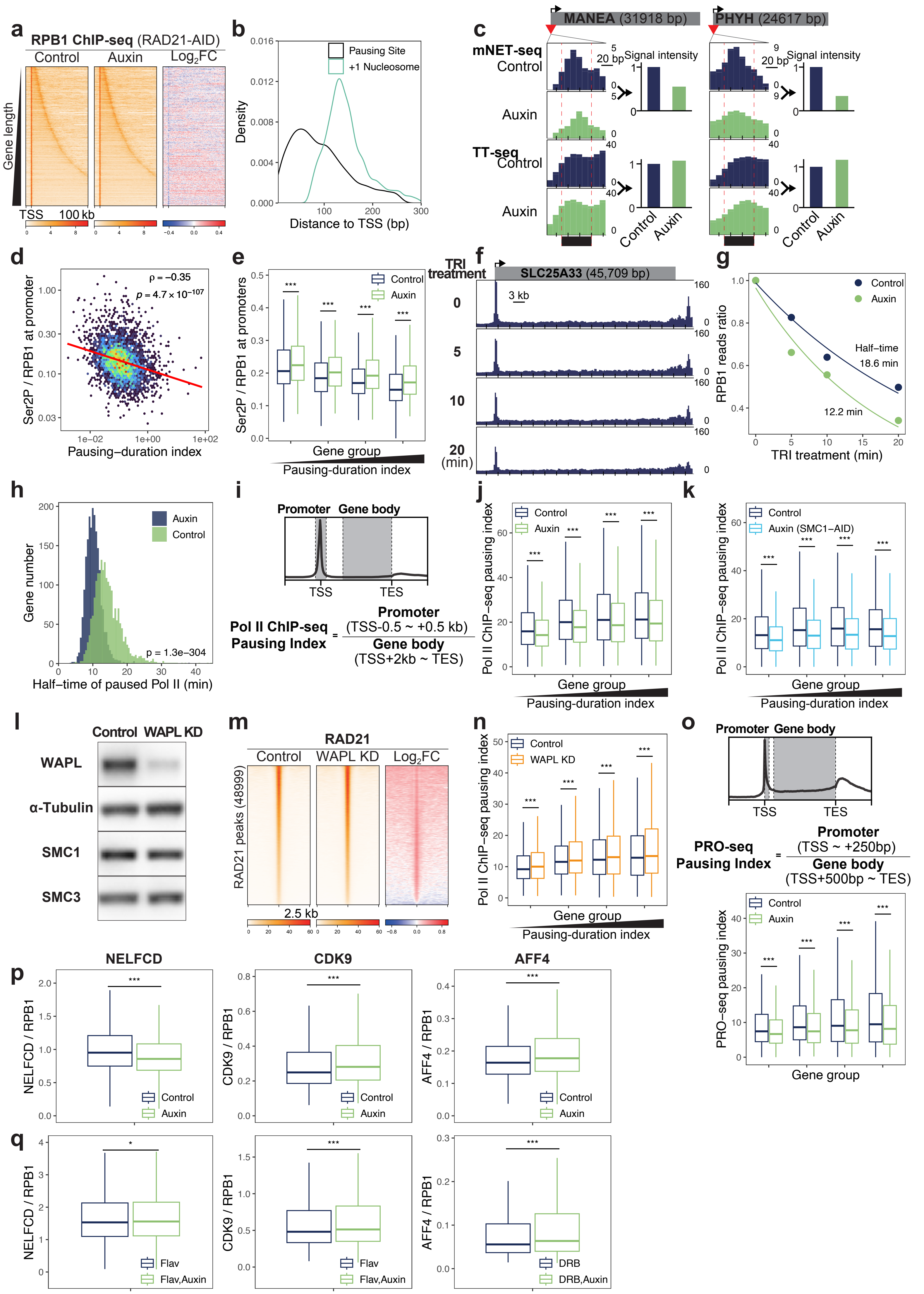
a
RPB1 ChIP-seq (RAD21-AID)
Control
Auxin
Log2FC
Gene length
TSS
100 kb
b
Pausing Site
+1 Nucleosome
Density
Distance to TSS (bp)
c
MANEA (31918 bp)
PHYH (24617 bp)
mNET-seq
Control
Auxin
TT-seq
Control
Auxin
20 bp
Signal intensity
Control
Auxin
d
ρ = −0.35
p = 4.7 × 10−107
Ser2P / RPB1 at promoter
Pausing−duration index
e
Control
Auxin
Ser2P / RPB1 at promoters
Gene group
Pausing-duration index
f
TRI treatment
SLC25A33 (45,709 bp)
3 kb
(min)
g
Control
Auxin
RPB1 reads ratio
Half-time
18.6 min
12.2 min
TRI treatment (min)
h
Auxin
Control
Gene number
p = 1.3e−304
Half−time of paused Pol II (min)
i
Promoter
Gene body
TSS
TES
Pol II ChIP-seq Pausing Index = Promoter (TSS-0.5 ~ +0.5 kb) / Gene body (TSS+2kb ~ TES)
j
Control
Auxin
Pol II ChIP−seq pausing index
Gene group
Pausing-duration index
k
Control
Auxin (SMC1−AID)
Pol II ChIP−seq pausing index
Gene group
Pausing-duration index
l
Control
WAPL KD
WAPL
α-Tubulin
SMC1
SMC3
m
RAD21
Control
WAPL KD
Log2FC
RAD21 peaks (48999)
2.5 kb
n
Control
WAPL KD
Pol II ChIP−seq pausing index
Gene group
Pausing-duration index
o
Promoter
Gene body
TSS
TES
PRO-seq Pausing Index = Promoter (TSS ~ +250bp) / Gene body (TSS+500bp ~ TES)
Control
Auxin
PRO−seq pausing index
Gene group
p
NELFCD
CDK9
AFF4
NELFCD / RPB1
CDK9 / RPB1
AFF4 / RPB1
Control
Auxin
q
NELFCD / RPB1
CDK9 / RPB1
AFF4 / RPB1
Flav
Flav,Auxin
DRB
DRB,Auxin

**Supplementary Fig. 3: Cohesin promotes Pol II pausing at highly paused genes.**
**a** Heatmaps showing RPB1 ChIP-seq signals across genes (TSS −10 kb to +100 kb) for 3,597 genes in control and auxin-treated cells. Genes are aligned at their transcription start site (TSS) and sorted by gene length.

**b** Histograms of distances from the TSSs to pausing sites and nucleosomes for 3,597 genes. Pausing sites were defined as the highest mNET-seq peak between the TSS and the +1 nucleosome. +1 nucleosome positions were determined by MNase–ChIP–seq (H3K4me3).

**c** Representative gene showing reduced mNET-seq signal at the pausing site upon cohesin depletion, while TT-seq signal at the same site remained largely unchanged. The red arrowhead indicates the position of the pausing site. Box plots showing quantified mNET-seq and TT-seq signal intensities within ±25 bp of the pausing site of the representative gene, used to calculate the pausing-duration index.

**d** Comparison of the pausing-duration index and Ser2P/RPB1 levels at promoters (TSS -1 kb to +1 kb) for 3,597 genes in control cells. P-value ($p$) and Spearman's correlation coefficient (ρ) were calculated using Spearman's rank correlation test. The red line indicates the linear regression fit

**e** Box plots showing Ser2P/RPB1 levels at promoters (TSS -1 kb to +1 kb) in control and auxin-treated cells. Gene groups are the same as those shown in Fig. 1d. Statistical significance was calculated using a two-sided Wilcoxon signed-rank test. (***$p < 0.001$).

**f** Representative gene showing reduced promoter-bound RNA polymerase II (Pol II) after triptolide (TRI) treatment. RPB1 ChIP-seq was performed at 0, 5, 10, and 20 min after TRI treatment.

**g** Exponential decay model estimating the half-life of paused Pol II at a representative gene shown in Supplementary Fig. 3e. RPB1 promoter occupancy (TSS −1 kb to +1 kb) was quantified at multiple time points after TRI treatment, and an exponential decay curve was fitted.

**h** Histograms of the half-life of paused Pol II in control and auxin-treated cells. Half-lives were estimated by fitting an exponential decay model to RPB1 promoter occupancy (TSS −1 kb to +1 kb) over time following triptolide (TRI) treatment. A total of 2,004 genes exhibiting a continual reduction in promoter RPB1 occupancy throughout the time-course experiment were analyzed. Mean half-lives: control, 14.4 min; auxin, 10.3 min. Statistical significance was calculated using a two-sided Wilcoxon signed-rank test.

**i** Pol II ChIP-seq pausing index is calculated as the ratio of Pol II occupancy at the promoter (TSS -0.5 kb to + 0.5 kb) to that in the gene body (TSS + 2kb to TES).

**j** Box plots showing Pol II ChIP-seq pausing index in control and auxin-treated cells. Gene groups are the same as those shown in Fig. 1d. Statistical significance was calculated using a two-sided Wilcoxon signed-rank test. (***$p < 0.001$). Exact p-values from left to right are: $3.80\times10^{-34}$, $1.36\times10^{-41}$, $4.32\times10^{-54}$, and $2.98\times10^{-27}$.

**k** Box plot showing Pol II ChIP-seq pausing index in control and auxin-treated cells expressing AID-tagged SMC1. Gene groups are the same as those shown in Fig. 1d. Statistical significance was calculated using a two-sided Wilcoxon signed-rank test. (***$p < 0.001$). Exact p-values from left to right are: $2.97\times10^{-75}$, $4.74\times10^{-72}$, $1.26\times10^{-64}$, and $8.63\times10^{-69}$.

**l** Western blots of WAPL, $\alpha$-tubulin, SMC1, and SMC3 in whole-cell extracts. Cells were treated with WAPL-targeting siRNA for two days. WAPL protein levels, normalized to $\alpha$-tubulin, decreased to ~22% of control levels, upon WAPL knockdown.

**m** Heatmaps showing RAD21 ChIP-seq siganls in control and WAPL-knockdown cells at 48,999 RAD21 ChIP-seq peaks. Peaks are sorted by RAD21 intensity in control cells.

**n** Box plots showing Pol II ChIP-seq pausing index in control and WAPL-knockdown cells. Gene groups are the same as those shown in Fig. 1d. Statistical significance was calculated using a two-sided Wilcoxon signed-rank test. (***$p < 0.001$). Exact p-values from left to right are: $1.37\times10^{-13}$, $1.27\times10^{-13}$, $1.39\times10^{-13}$, and $1.74\times10^{-16}$.

**o** PRO-seq pausing index is calculated as the ratio of PRO-seq signal density at the promoter (TSS to +250 bp) to that in the gene body (TSS + 500 bp to TES). Box plots of PRO-seq pausing index in control and auxin-treated cells. PRO-seq data were from Rao et al. (2017). Gene groups are the same as those shown in Fig. 1d. Statistical significance was calculated using a two-sided Wilcoxon signed-rank test. (***$p < 0.001$). Exact p-values from left to right are: $4.58\times10^{-42}$, $1.94\times10^{-50}$, $4.54\times10^{-46}$, and $2.19\times10^{-42}$.

**p** Box plots showing the relative levels of NELFCD, CDK9, and AFF4 to RPB1 promoters (TSS -1 kb to +1 kb) for 3,597 genes in control and auxin-treated cells. Statistical significance was calculated using a two-sided Wilcoxon signed-rank test. (***$p < 0.001$). Exact p-values are: $1.89\times10^{-55}$ (NELFCD), $2.07\times10^{-58}$ (CDK9), and $1.30\times10^{-35}$ (AFF4).

**q** Box plots showing the relative levels of NELFCD, CDK9, and AFF4 to RPB1 at promoters (TSS -1 kb to +1 kb) for 3,597 genes in control and auxin-treated cells, in the presence of flavopiridol or DRB. Statistical significance was calculated using a two-sided Wilcoxon signed-rank test. (*$p < 0.05$, ***$p < 0.001$). Exact p-values are: $4.00\times10^{-2}$ (NELFCD), $1.26\times10^{-15}$ (CDK9), and $2.05\times10^{-77}$ (AFF4).

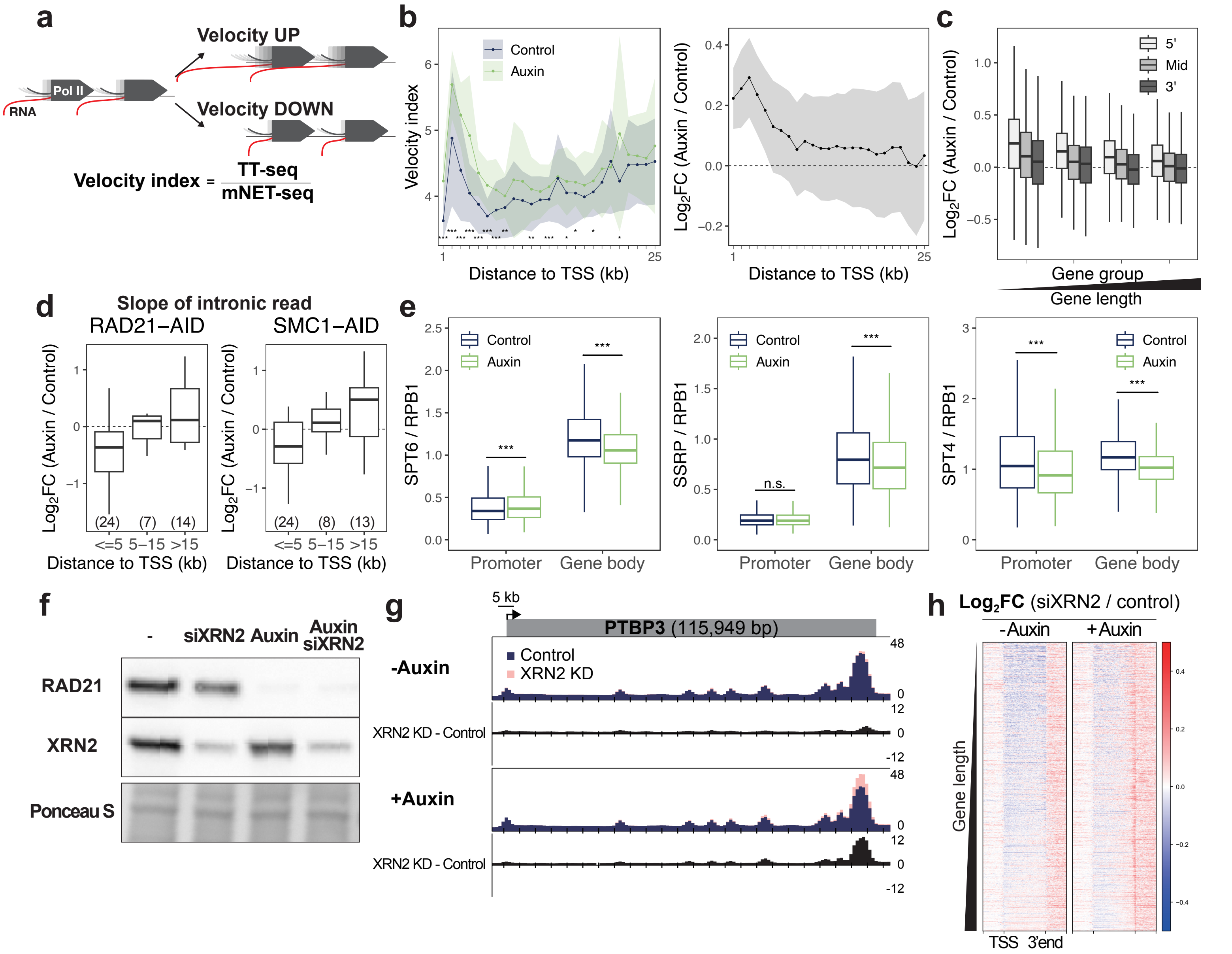


**Supplementary Fig. 4: Cohesin constrains early Pol II elongation and stabilizes transcription elongation to prevent premature termination.**

**a** Velocity index was calculated as the ratio of TT-seq to mNET-seq across gene bodies.

**b** Line plots with error shading showing the velocity index in control and auxin-treated cells (left), and their $\log_2$ fold changes (auxin/control) (right). Values are shown from the TSS to +25 kb in 1-kb bins, where available, using data from 3,597 genes. Data represent mean ± 95% confidence intervals. Statistical significance was calculated using a two-sided Wilcoxon signed-rank test. (**p* < 0.05, ***p* < 0.01, ****p* < 0.001).

**c** Box plots of $\log_2$ fold changes (auxin/control) in the velocity index across gene bodies. Gene bodies were divided into three equal regions (5', Mid, and 3') from the TSS to 3'gene end. A total of 3,597 genes were evenly divided into four groups based on their length.

**d** Box plots showing $\log_2$ fold changes (auxin/control) in the slope of RNA-seq intronic reads in RAD21-depleted (left) and SMC1-depleted cells (right). The slope was estimated by fitting a linear model to intronic read coverage. Introns longer than 1 kb with a negative slope and an adjusted $R^2 > 0.5$ were included in the analysis. The number of introns analyzed is indicated in parentheses.

**e** Box plots of the relative levels of SPT6 (left), SSRP (FACT complex) (mid), and SPT4 (DSIF complex) (right) to RPB1 in control and auxin-treated cells at promoters (TSS -1 kb to +1 kb) and gene bodies (TSS +1 kb to 3'gene end) for 3,597 genes. Statistical significance was calculated using a two-sided Wilcoxon signed-rank test. (****p* < 0.001). Exact p-values: $1.78\times10^{-26}$ (SPT6 promoter), $8.75\times10^{-249}$ (SPT6 gene body), 0.428 (SSRP promoter), $3.91\times10^{-145}$ (SSRP gene body), $4.48\times10^{-69}$ (SPT4 promoter), $<1\times10^{-300}$ (SPT4 gene body).

**f** Western blots of RAD21 and XRN2, with Ponceau staining in whole-cell extract. XRN2 protein levels, normalized to Ponceau, decreased to ~22% of control levels upon siXRN2 treatment.

**g** Representative gene showing increased RNA-seq signal upon XRN2 knockdown, which was further enhanced upon auxin-induced cohesin depletion. Difference tracks (XRN2 KD – Control) are shown below each condition.

**h** Heatmaps of $\log_2$ fold changes (siXRN2/control and auxin + siXRN2/auxin) in RNA-seq across genes (TSS to 3' gene end) for 3,597 genes. Genes are aligned at their 3'gene end and sorted by gene length.

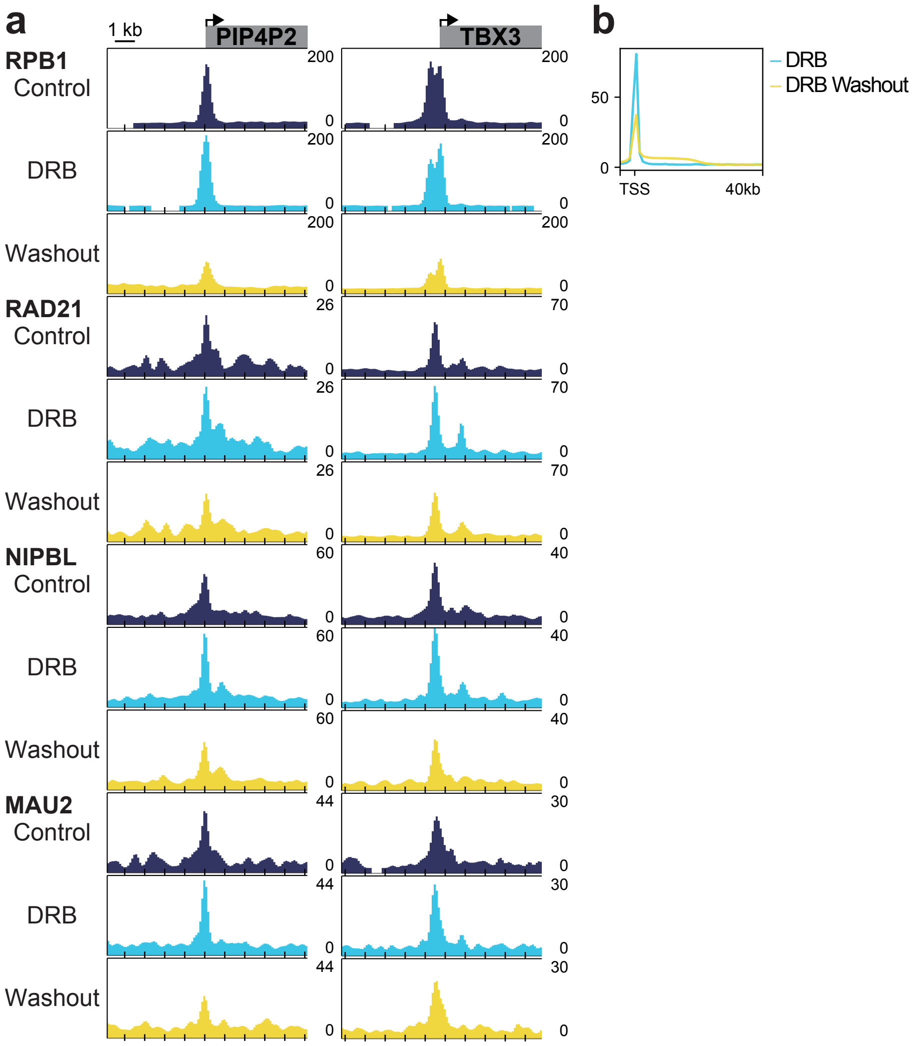


**Supplementary Fig. 5: Cohesin and its loader coordinatedly colocalize with RNA Polymerase II at promoters.**
**a** Representative genes showing increased promoter binding of RPB1, RAD21, NIPBL, and MAU2 upon DRB treatment, which was reduced upon DRB washout (TSS –5 kb to +5 kb).

**b** Average ChIP-seq profiles of RPB1 under DRB treatment and DRB washout conditions at 1,766 genes longer than 40 kb (TSS -5 kb to +40 kb).

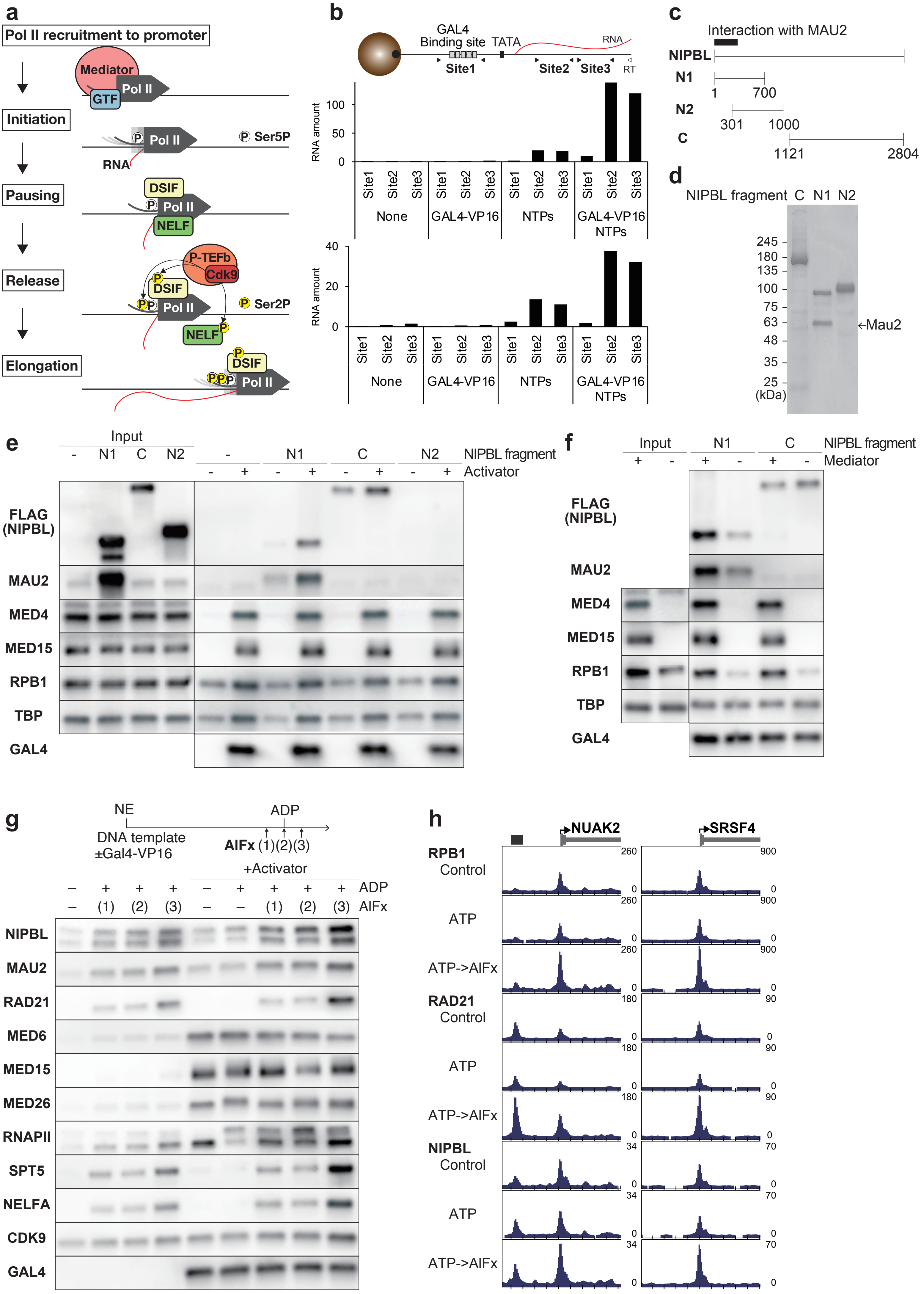

a
Pol II recruitment to promoter
Mediator
Pol II
GTF
Initiation
Pol II
Ser5P
RNA
Pausing
DSIF
Pol II
NELF
Release
P-TEFb
Cdk9
DSIF
Pol II
Ser2P
NELF
Elongation
DSIF
Pol II
b
GAL4
Binding site
TATA
RNA
Site1
Site2
Site3
RT
RNA amount
Site1
Site2
Site3
None
GAL4-VP16
NTPs
GAL4-VP16 NTPs
c
Interaction with MAU2
NIPBL
N1
1
700
N2
301
1000
C
1121
2804
d
NIPBL fragment C N1 N2
245
180
135
100
75
63
48
35
25
(kDa)
←Mau2
e
Input
- N1 C N2
- N1 C N2 NIPBL fragment
- + - + - + - + Activator
FLAG (NIPBL)
MAU2
MED4
MED15
RPB1
TBP
GAL4
f
Input N1 C NIPBL fragment
+ - + - + - Mediator
FLAG (NIPBL)
MAU2
MED4
MED15
RPB1
TBP
GAL4
g
NE
DNA template ±Gal4-VP16
ADP
AlFx (1)(2)(3)
+Activator
– + + + – + + + + ADP
– (1) (2) (3) – – (1) (2) (3) AlFx
NIPBL
MAU2
RAD21
MED6
MED15
MED26
RNAPII
SPT5
NELFA
CDK9
GAL4
h
NUAK2
SRSF4
RPB1
Control
ATP
ATP->AlFx
RAD21
Control
ATP
ATP->AlFx
NIPBL
Control
ATP
ATP->AlFx
260
900
180
90
34
70
0

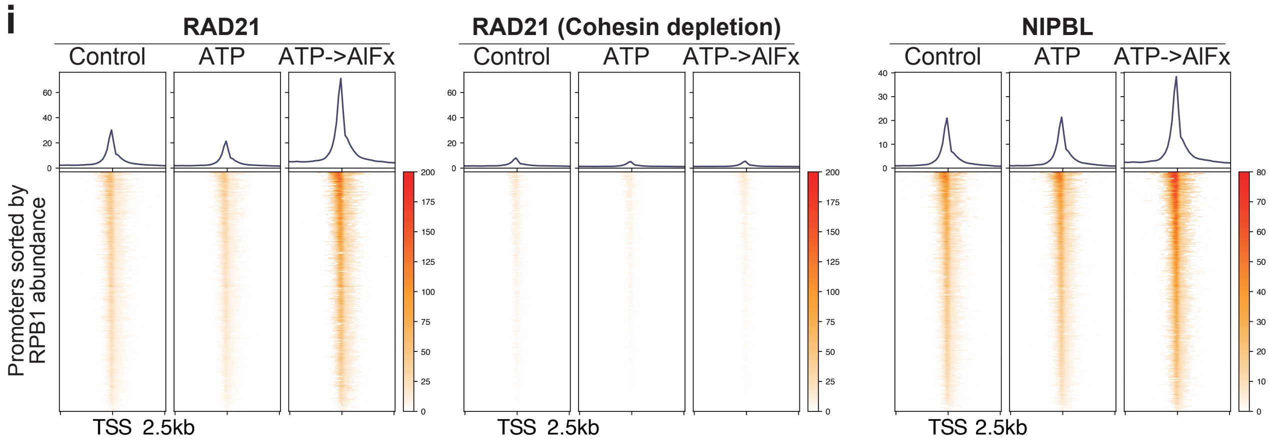


**Supplementary Fig. 6: Cohesin and its loader engage the transcription machinery at the pause–release transition.**
**a** Schematic illustration of transcriptional stages. First, RNA polymerase II (Pol II) is recruited to promoters bound by general transcription factors (GTFs) and the Mediator complex, and it initiates transcription. Shortly after initiation, Pol II pauses with the help of NELF and DSIF. CDK9, a component of P-TEFb, phosphorylates the Pol II C-terminal domain (CTD), NELF, and DSIF to promote pause release. Pol II then enters productive elongation.

**b** Schematic of primer design to detect RNA production (top). For qPCR, Site1 targets the GAL4-binding sites, whereas Site2 and Site3 target the gene body downstream of the TATA box. RT indicates the position of the oligonucleotide used for reverse transcription. Box plots show RNA amounts quantified by qPCR under the following conditions: none, GAL4-VP16, NTPs, and GAL4-VP16 + NTPs, from two independent replicates (bottom). A value of 1 corresponds to the signal equivalent to 1 μg of template DNA as determined by a standard curve of the template DNA.

**c** Schematic representation of the three recombinant NIPBL fragments used in the *in vitro* transcription assay. The black box indicates the N-terminal region of NIPBL required for interaction with MAU2.

**d** Coomassie blue staining of purified recmbinant NIPBL fragments (C, N1, and N2). The fragments were expressed in insect cells and purified. MAU2 was co-expressed with the N1 fragment and co-purified.

**e** *In vitro* transcription assay with NIPBL fragments. Nuclear extracts (Input) were mixed with FLAG-tagged recombinant NIPBL fragments, then incubated with template DNA preloaded with GAL4-VP16.

**f** *In vitro* transcription assay with Mediator depletion on the recruitment of NIPBL mutants (N1 and C). The mediator complex was depleted from nuclear extract using antibodies against MED4, MED15, and MED25. Nuclear extracts (Input) were mixed with FLAG-tagged recombinant NIPBL fragments, then incubated with template DNA preloaded with GAL4-VP16.

**g** *In vitro* transcription assay with cohesin ATPase inhibition by AlFx. Template DNA, preloaded with GAL4-VP16, was incubated with nuclear extracts (Input). ADP was added 20 min after the start of incubation. AlFx was added either 3 minutes before (1), simultaneously with (2), or 3 minutes after (3) ADP addition.

**h** Representative gene showing enhanced promoter binding of RPB1, RAD21, and NIPBL upon ATP->AlFx treatment. Black box indicates the position of a cohesin peak.

**i** Heatmaps of RAD21 and NIPBL CUT&Tag signals at promoters (TSS −2.5 kb to +2.5 kb) for 3,597 genes under control, ATP, and ATP->AlFx conditions, with or without cohesin depletion. Permeabilized cells were treated with ATP for 10 min or with ATP for 5 min followed by AlFx for 5 min.

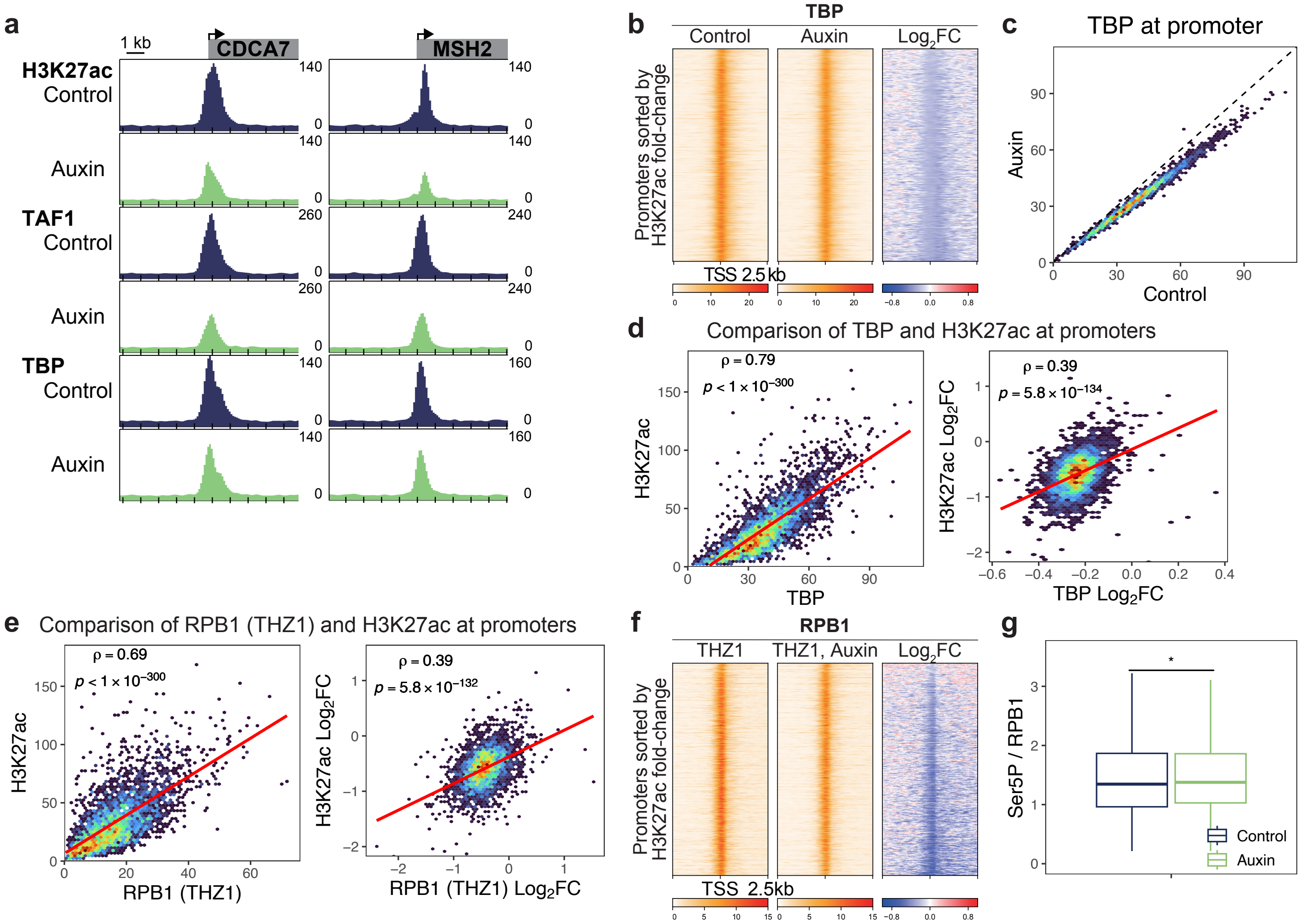


**Supplementary Fig. 7: Cohesin maintains the H3K27ac mark at promoters.**
**a** Representative genes showing decreased promoter binding of H3K27ac, TAF1, and TBP upon auxin treatment (TSS –5 kb to +5 kb).

**b** Heatmaps of TBP ChIP-seq signals at promoters (TSS -2.5 kb to +2.5 kb) in control and auxin-treated cells. Promoters are shown in the same order as in Fig. 5a.

**c** Comparison of promoter TBP (TSS -1 kb to +1 kb) between control and auxin-treated cells for 3,597 genes. Upon cohesin depletion, TBP levels decreased at 92% of promoters and increased at 8% (FDR < 0.05, binomial test).

**d** Comparison between promoter TBP and H3K27ac (TSS -1 kb to +1 kb) in control cells, and their $\log_2$ fold changes (auxin/control) for 3,597 genes. P-value (*p*) and Spearman's correlation coefficient (ρ) were calculated using Spearman's rank correlation test.

**e** Heatmaps of RPB1 ChIP-seq signals in the presence of THZ1 at promoters (TSS -2.5 kb to +2.5 kb) in control and auxin-treated cells. Promoters are shown in the same order as in Fig. 5a.

**f** Comparison between promoter RPB1 in the presence of THZ1 and H3K27ac (TSS -1 kb to +1 kb) in control cells (left), and their $\log_2$ fold changes (auxin/control) for 3,597 genes (right). P-value (*p*) and Spearman's correlation coefficient (ρ) were calculated using Spearman's rank correlation test.

**g** Box plots of the relative Ser5P levels to RPB1 at promoters (TSS -1 kb to +1 kb) for 3,597 genes in control and auxin-treated cells. Statistical significance was calculated using a two-sided Wilcoxon signed-rank test. (*$p < 0.05$). Exact p-values is: $1.67 \times 10^{-2}$.

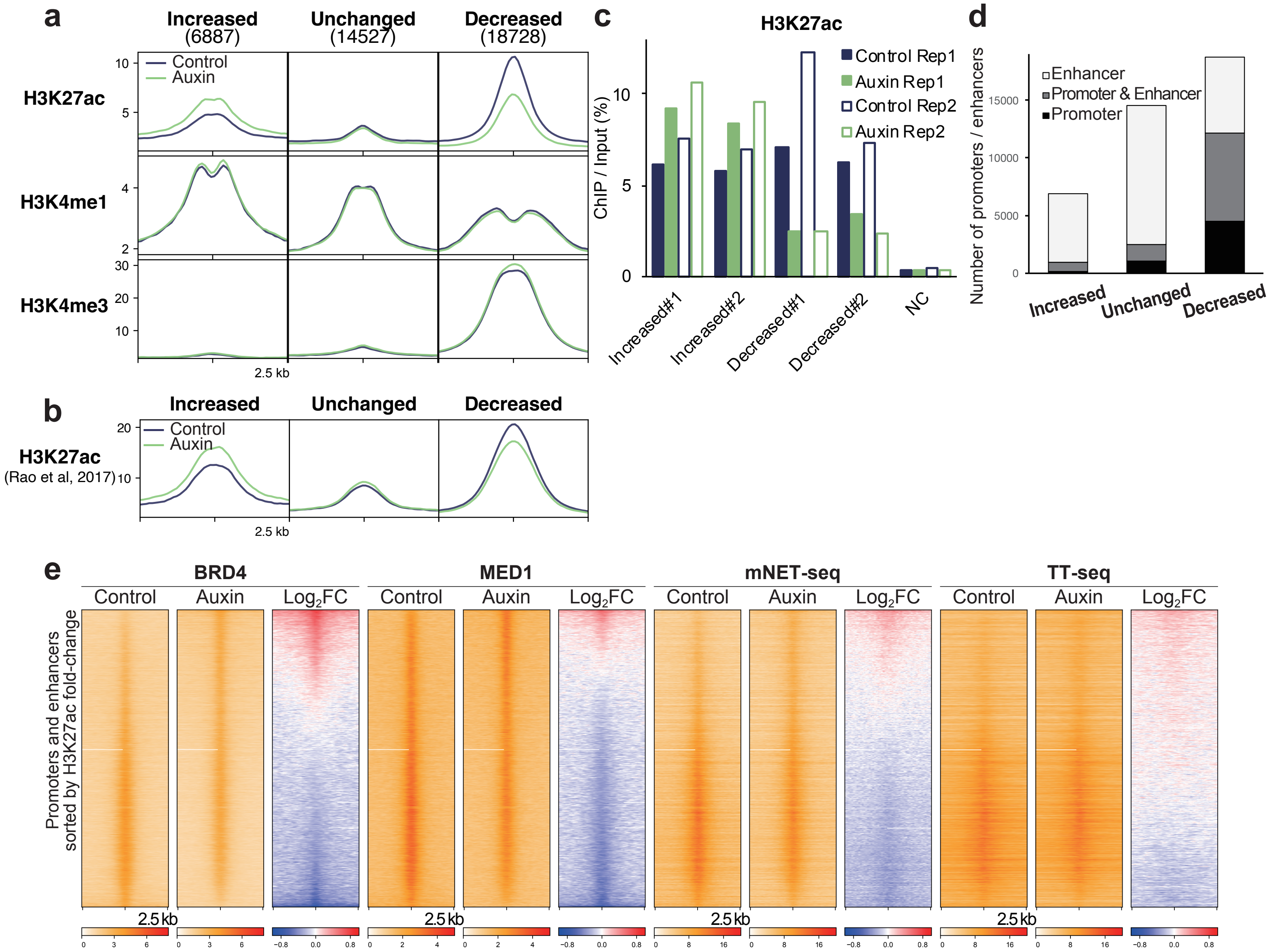


**Supplementary Fig. 8: Cohesin regulates H3K27ac distribution via promoter–enhancer contacts.**

**a** Average ChIP-seq profiles of H3K27ac, H3K4me1, and H3K4me3 at regulatory elements (promoters and enhancers). Regulatory elements were classified as increased, unchanged, or decreased based on their H3K27ac response to cohesin depletion. The number of elements in each category is shown in parentheses.

**b** Average ChIP-seq profiles of H3K27ac from Rao et al. (2017).at regulatory elements (promoters and enhancers). Regulatory elements were classified as increased, unchanged, or decreased as in a.

**c** ChIP–qPCR of H3K27ac at four representative regulatory elements and a negative control (NC) site. Two elements were selected from regions with increased H3K27ac and two from regions with decreased H3K27ac, based on ChIP–seq results. Data are shown for two biological replicates (Rep1 and Rep2).

**d** Bar plots of the number of regulatory elements annotated as promoter or enhancer by ChromHMM. Regulatory elements were categorized as in Fig. 6e (increased, unchanged, or decreased).

**e** Heatmaps of BRD4 and MED1 ChIP-seq, alongside mNET-seq and TT-seq signals in control and auxin-treated cells at regulatory elements. Regulatory elements are shown in the same order as in Fig. 6a.

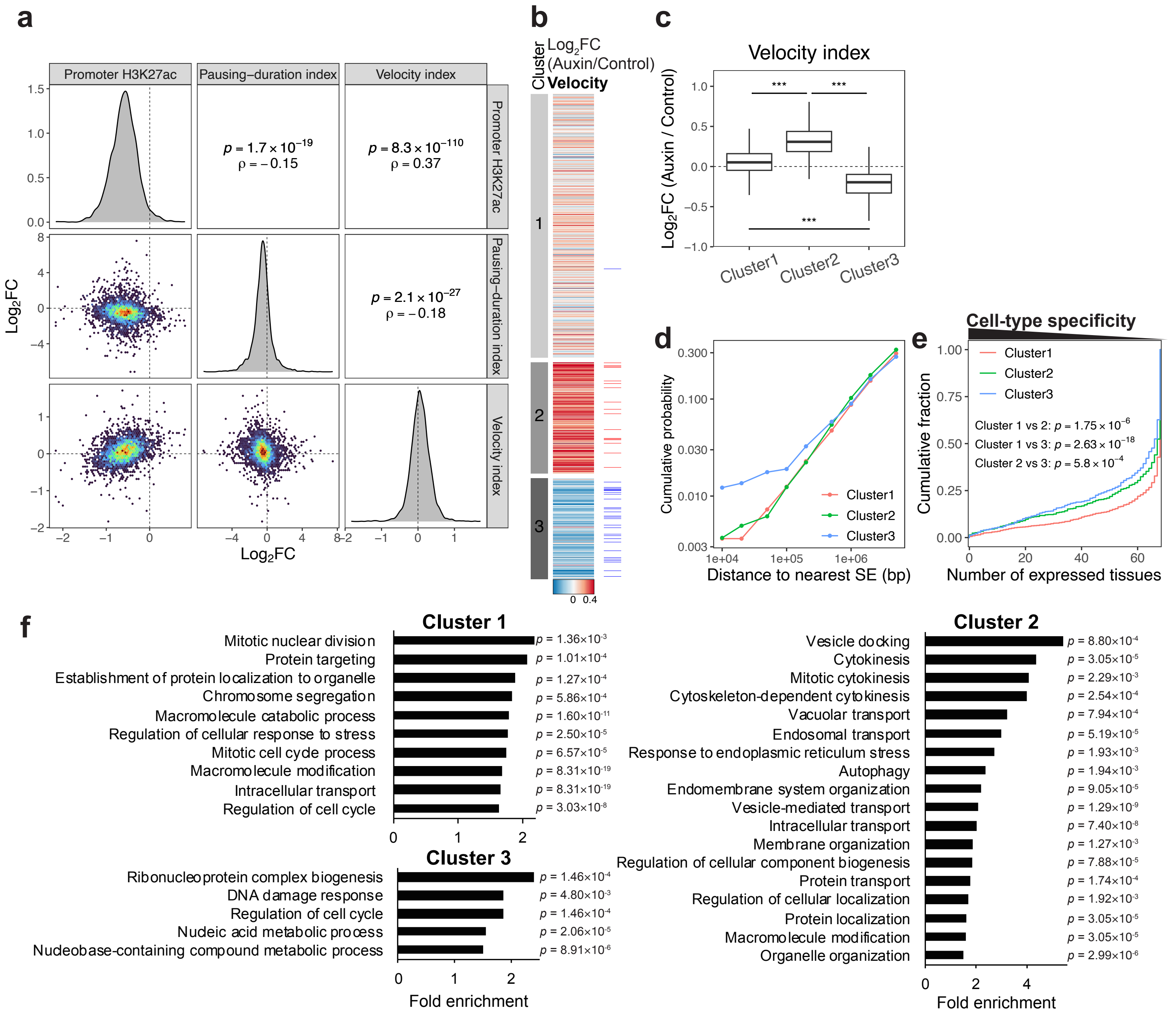


**Supplementary Fig. 9: Genes with modest expression changes upon cohesin loss show reduced Pol II recruitment and enhanced pause release.**

**a** Pair plots showing the relationships among $\log_2$ fold changes in promoter H3K27ac, pausing-duration index, and velocity index. Velocity index was calculated as the TT-seq/mNET-seq ratio across gene bodies (TSS +1 kb to 3'end –1 kb). Diagonal panels show the distribution of each variable. Lower triangular panels show pairwise comparisons, and upper triangular panels display P-value ($p$) and Spearman's correlation coefficient ($\rho$) from Spearman's rank correlation test.

**b** Heatmaps showing $\log_2$ fold changes (auxin/control) in velocity index for each gene. Genes are shown in the same order and cluster assignment as in Fig. 7a. Genes identified as upregulated or downregulated by TT-seq (Fig. 1a) are highlighted with red and blue lines, respectively.

**c** Box plots showing $\log_2$ fold changes (auxin/control) in the velocity index across Clusters 1, 2, and 3 shown in Fig. 7a. Statistical significance was calculated using multiple pairwise Welch's t-tests between gene groups. (***$p < 0.001$). Exact p-values for velocity index: $2.13\times10^{-190}$ (Cluster1 vs 2), $2.89\times10^{-192}$ (Cluster1 vs 3), $3.41\times10^{-232}$ (Cluster2 vs 3).

**d** Cumulative probability distributions of distances to the nearest super enhancers for genes clustered into Cluster 1–3.

**e** Cumulative distributions of cell-type specificity for genes clustered into Cluster 1–3. Statistical significance was calculated using the Kolmogorov–Smirnov test.

**f** GO term analysis of genes clustered into Cluster 1–3. Only GO terms with a fold enrichment greater than 1.5 and a Benjamini-adjusted P value below 0.05 were considered significant and are shown.

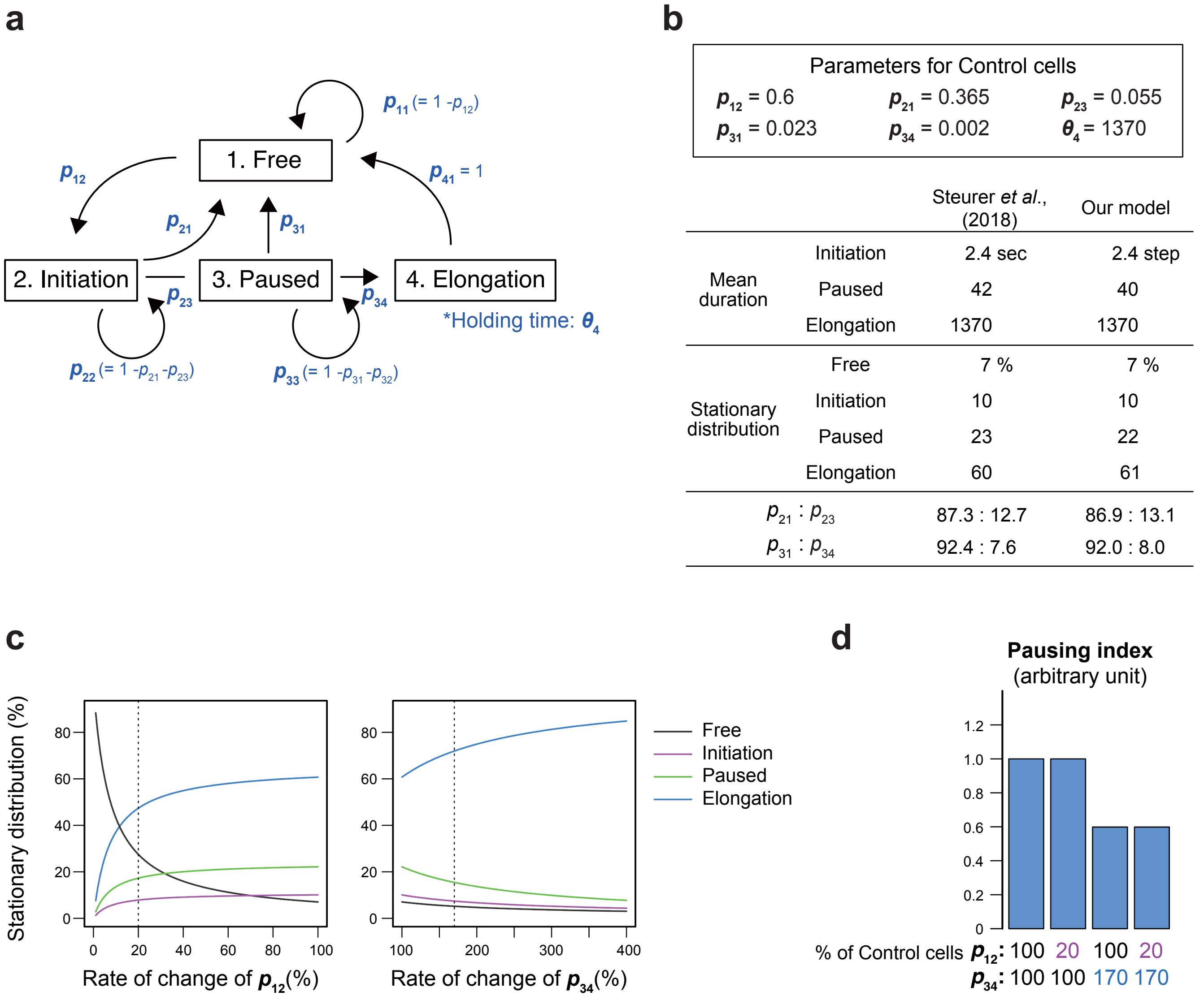


**Supplementary Fig. 10: Simulations suggest compensatory effects of reduced Pol II recruitment and enhanced pause release.**

**a** Schematic of the transcriptional states and transitions in our model. Pol II moves through four states: Free (1), Initiation (2), Paused (3), and Elongation (4). Transitions between these states are governed by probabilities $p_{ij}$, where $p_{ij}$ denotes the probability of transition from state i to state j. The holding time for the Elongation state is denoted by $\theta_4$. The model includes six independent parameters.

**b** Estimated parameters and model validation. (Upper) Optimized parameters that best matched the intracellular Pol II dynamics. (Lower) Comparison of key indicators of Pol II dynamics between experimentally inferred values (Steurer et al., 2018) and our model's predictions using these optimized parameters.

**c** Line plots showing the impact of varying $p_{12}$ (0-100%) and $p_{34}$ (100-400%) on the stationary distribution. The vertical dotted line indicates the parameter set used in Figure S9D, representing the condition under cohesin depletion.

**d** Bar plots showing simulated effects of cohesin depletion on the pausing index. Our experimental data revealed that cohesin depletion reduces transcription initiation and enhances pause release. In our model, these changes correspond to decreased $p_{12}$ and increased $p_{34}$, and each parameter is altered individually or in combination. The pausing index, designed to mimic the Pol II ChIP-seq pausing index, was calculated as ([I] + [P])/[E], where [X] represents the probability of Pol II being in state X. The values were normalized to 1 for the control condition.

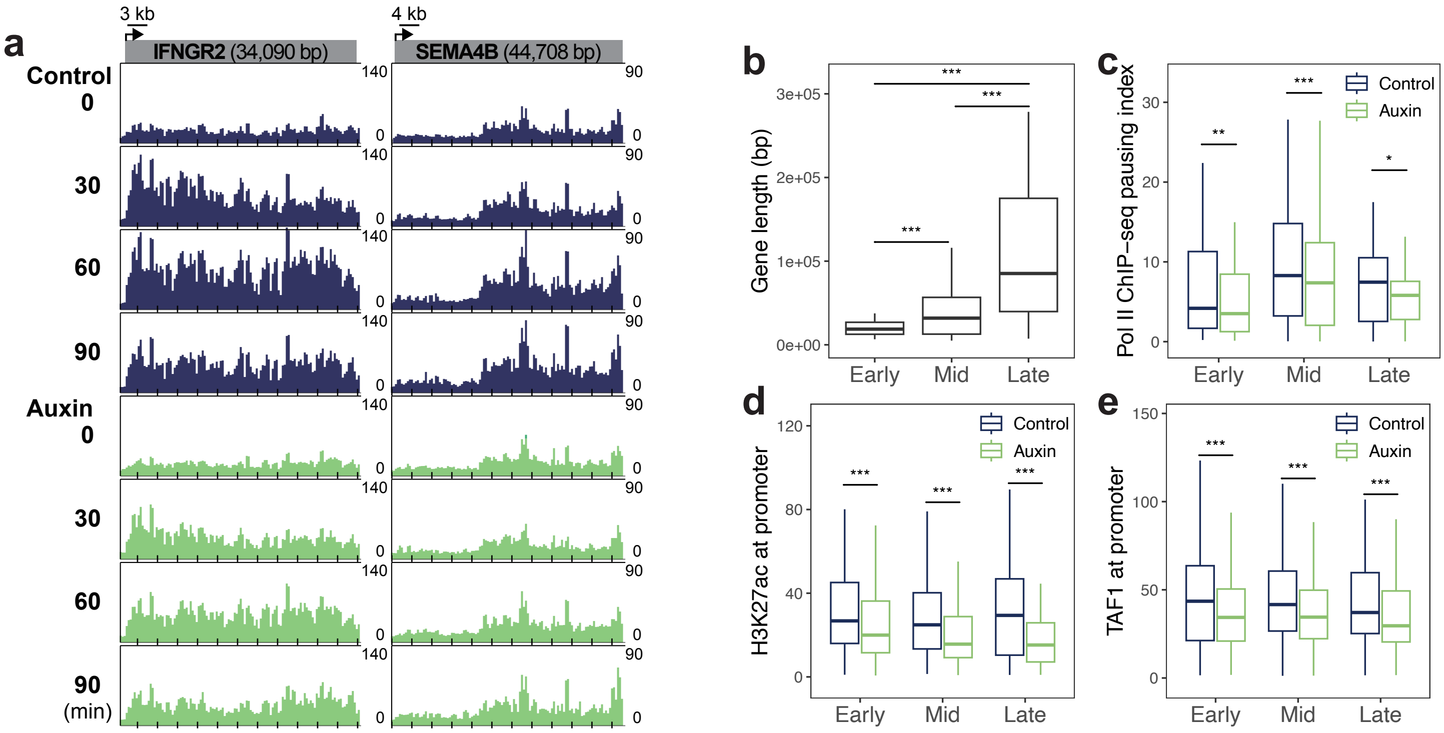


**Supplementary Fig. 11: Cohesin depletion impairs rapid gene induction.**

**a** Representative genes showing TNFα-induced expression that was reduced by auxin treatment. EU-seq was performed in control and auxin-treated cells at 0, 30, 60, and 90 min after TNFα treatment.

**b** Box plots showing gene lengths of early-, mid-, and late-induced genes identified in Fig. 7e. Statistical significance was calculated using multiple pairwise Welch's t-tests between gene groups. (***$p$ < 0.001).

**c** Box plots showing Pol II ChIP-seq pausing index for early-, mid-, and late-induced genes in control and auxin-treated cells. Statistical significance was calculated using a two-sided Wilcoxon signed-rank test. (*$p$ < 0.05, **$p$ < 0.01, ***$p$ < 0.001).

**d** Box plots showing promoter H3K27ac (TSS -1 kb to +1 kb) for early-, mid-, and late-induced genes in control and auxin-treated cells. Statistical significance was calculated using a two-sided Wilcoxon signed-rank test. (***$p$ < 0.001).

**e** Box plots showing promoter TAF1 (TSS -1 kb to +1 kb) at early-, mid-, and late-induced genes in control and auxin-treated cells. Statistical significance was calculated using a two-sided Wilcoxon signed-rank test. (***$p$ < 0.001).

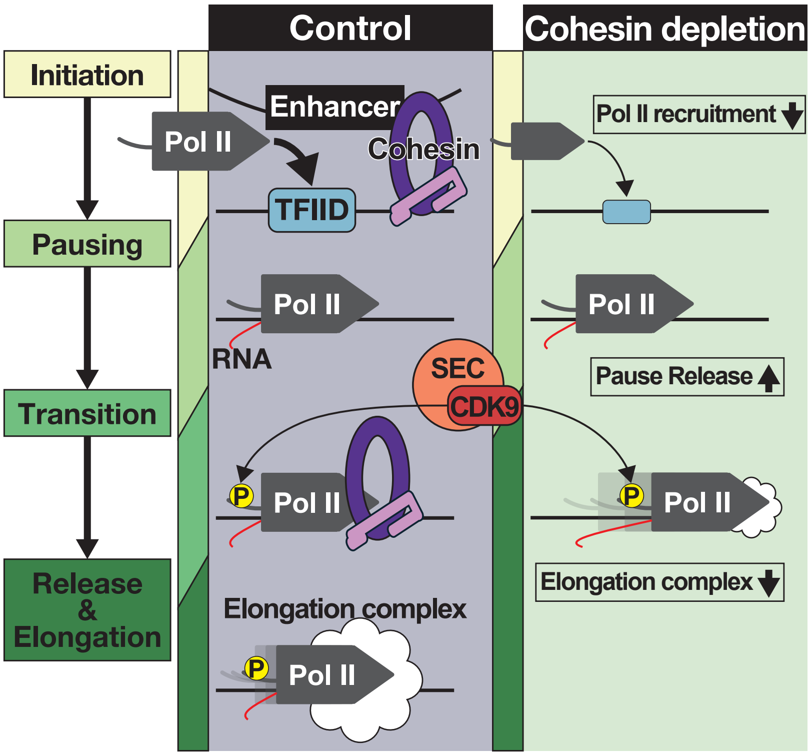


**Supplementary Fig. 12: Schematic model showing how cohesin regulates transcription.**
Cohesin promotes Pol II recruitment by facilitating promoter–enhancer communication. It also restrains pause release by engaging the transcriptional machinery at the pausing-to-release transition. This engagement functions as a quality-control-like step that coordinates elongation factor recruitment and prevents premature transcription termination. Upon cohesin depletion, Pol II recruitment decreases while pause release is enhanced. These compensatory effects contribute to the minimal changes in steady-state gene expression. However, shortened pausing duration leads to immature formation of the elongation complex and premature transcription termination.

**Supplementary Table 1. Spearman correlation between biological replicates**

| Method | Condition | Comparison | Spearman correlation |
|---|---|---|---|
| TT-seq | Control | Rep1 vs Rep2 | 0.812770766 |
| | Auxin | Rep1 vs Rep2 | 0.761359918 |
| RNA-seq | Control | Rep1 vs Rep2 | 0.96174539 |
| | | Rep1 vs Rep3 | 0.956172489 |
| | | Rep2 vs Rep3 | 0.956834384 |
| | Auxin | Rep1 vs Rep2 | 0.959113347 |
| | | Rep1 vs Rep3 | 0.953740064 |
| | | Rep2 vs Rep3 | 0.955085477 |
| | siXRN2 | Rep1 vs Rep2 | 0.961018929 |
| | | Rep1 vs Rep3 | 0.954608991 |
| | | Rep2 vs Rep3 | 0.95577589 |
| | siXRN2+Auxin | Rep1 vs Rep2 | 0.960277533 |
| | | Rep1 vs Rep3 | 0.954370695 |
| | | Rep2 vs Rep3 | 0.954522163 |
| | Control (SMC1-AID) | Rep1 vs Rep2 | 0.957102357 |
| | | Rep1 vs Rep3 | 0.955258042 |
| | | Rep2 vs Rep3 | 0.954701116 |
| | Auxin (SMC1-AID) | Rep1 vs Rep2 | 0.961098241 |
| | | Rep1 vs Rep3 | 0.956859115 |
| | | Rep2 vs Rep3 | 0.955936042 |
| EU-seq | Control | Rep1 vs Rep2 | 0.946378543 |
| | Auxin | Rep1 vs Rep2 | 0.949427226 |
| ChIP-seq (RPB1) | Control | Rep1 vs Rep2 | 0.9194972 |
| | Auxin | Rep1 vs Rep2 | 0.930044655 |
| mNET-seq | Control | Rep1 vs Rep2 | 0.766972871 |
| | | Rep1 vs Rep3 | 0.794196871 |
| | | Rep2 vs Rep3 | 0.783041907 |
| | Auxin | Rep1 vs Rep2 | 0.74417042 |
| | | Rep1 vs Rep3 | 0.768983191 |
| | | Rep2 vs Rep3 | 0.785839327 |
| ChIP-seq (NELFCD) | Control | Rep1 vs Rep2 | 0.81995541 |
| | Auxin | Rep1 vs Rep2 | 0.812821833 |
| ChIP-seq (CDK9) | Control | Rep1 vs Rep2 | 0.903349594 |
| | Auxin | Rep1 vs Rep2 | 0.898996258 |
| ChIP-seq (AFF4) | Control | Rep1 vs Rep2 | 0.853833068 |
| | Auxin | Rep1 vs Rep2 | 0.845125655 |
| ChIP-seq (H3K27ac) | Control | Rep1 vs Rep2 | 0.91940095 |
| | Auxin | Rep1 vs Rep2 | 0.917521016 |
| ChIP-seq (TBP) | Control | Rep1 vs Rep2 | 0.944027499 |
| | Auxin | Rep1 vs Rep2 | 0.917486999 |
| ChIP-seq (PAF) | Control | Rep1 vs Rep2 | 0.906040642 |
| | Auxin | Rep1 vs Rep2 | 0.892281724 |
| ChIP-seq (SPT6) | Control | Rep1 vs Rep2 | 0.886966271 |
| | Auxin | Rep1 vs Rep2 | 0.886047038 |

**Supplementary Table 2. QuASAR-Rep reproducibility scores for Micro-C replicates**

| Method | Condition | Comparison | QuASAR-Rep reproducibility score |
|---|---|---|---|
| Micro-C | Control | Rep1 vs Rep2 | 0.86512 |
| | Auxin | Rep1 vs Rep2 | 0.875007 |
